# Learning to cooperate: The evolution of social rewards in repeated interactions

**DOI:** 10.1101/074096

**Authors:** Slimane Dridi, Erol Akçay

## Abstract

Understanding the behavioral and psychological mechanisms underlying social behaviors is one of the major goals of social evolutionary theory. In particular, a persistent question about animal cooperation is to what extent it is supported by other-regarding preferences. In many situations, animals adjust their behaviors through learning by responding to the rewards they experience as a consequence of their actions. Therefore, we may ask whether learning in social situations can be driven by evolved prosocial rewards. Here we develop a mathematical model in order to ask whether the mere act of cooperating with a social partner will evolve to be inherently rewarding. Individuals interact repeatedly in pairs and adjust their behaviors through reinforcement learning. We assume that individuals associate to each game outcome an internal reward value. These perceived rewards are genetically evolving traits. We find that conditionally cooperative rewards that value mutual cooperation positively but the sucker’s outcome negatively tend to be evolutionarily stable. Purely other-regarding rewards can evolve only under special parameter combinations. On the other hand, selfish rewards that always learn pure defection are also evolutionary successful. These findings are consistent with empirical observations showing that humans tend to show conditionally cooperative behavior, and also exhibit diversity of preferences. Our model also demonstrates the need to further integrate multiple levels of biological causation of behavior.

## Introduction

In animals, repeated interactions often lead to mutual cooperation (Trivers 1971; Axelrod and Hamilton 1981; Wilkinson 1988; Lehmann and Keller 2006; Schneeberger et al. 2012; Stewart and Plotkin 2013). Because repeated interactions offer the opportunity for learning, there is growing interest in characterizing the learning mechanisms and internal social motivations that lead to cooperation. Recognizing that natural selection acts on those behavioral mechanisms (McNamara and Houston 2009; Hammerstein and Stevens 2012; Fawcett et al. 2013; Dridi and Lehmann 2014) rather than directly on the cooperative phenotypes themselves generates a new perspective on questions about the evolution of cooperation. In particular, an important question at the interface of psychological mechanisms and evolutionary theory is whether biological altruism requires or necessarily leads to psychological altruism, also called other-regarding preferences. In other words, when we observe cooperation, is it because the individuals performing the cooperative act have other-regarding preferences, i.e., they evolved motivations to provide a positive outcome for their social partners? This question about the proximate mechanisms underlying cooperation is important both to understand how individuals will behave in novel social and environmental contexts but also how natural selection will shape the evolution of social traits (Akçay et al. 2009; Akçay and Van Cleve 2012; Van Cleve and Akçay 2014).

Several studies of cooperation in animals suggest that individuals may have other-regarding preferences (mostly in primates; Brosnan et al. 2010; Chang et al. 2011; Claidière et al. 2015; Lakshminarayanan and Santos 2008, but recently also in rats Hernandez-Lallement et al. 2015). However, other studies found that animals seem to pursue only their own personal gain (Jensen et al. 2006; Silk et al. 2005). In these experiments, animals are generally presented with the choice between a selfish option (obtaining a reward only for oneself) and a social option (providing a reward for both oneself and a partner), and a preference for the social option may be interpreted as other-regarding preferences. Evidence for such prosocial tendencies is also abundant in humans (Fehr and Gächter 2000; Henrich et al. 2001; Camerer 2003; Fehr and Fischbacher 2003; Chaudhuri 2010). Nonetheless, other researchers (Binmore 2005; Burton-Chellew et al. 2015, 2016) argue that apparently other-regarding behavior may be explained by the participants not fully understanding the experiment’s setup, combined with payoff-based learning during the course of the game.

A much greater number of empirical studies (Taborsky et al. 2016) indicate that reciprocal cooperation occurs in species as diverse as fish (Dugatkin and Alfieri 1991), birds (Voelkl et al. 2015), bats (Wilkinson et al. 2016), or primates (Schino and Aureli 2010). However, our understanding of the psychological motives underlying reciprocation remains limited. In particular, reciprocal cooperation can come about from simple reaction norms (McNamara et al. 1999), other-regarding preferences (Akçay et al. 2009), or learning from past rewards. In short, there is still disagreement about whether prosocial preferences combined with learning explain the cooperation that we observe in human and other animal societies. Answering this question requires explicit models that combines behavioral dynamics of learning with evolution of rewards, which is what motivates this paper.

Much of standard economics and decision theory is built on the idea that individuals strive to maximize a quantity called “utility.” Likewise, one can show that in the long run, natural selection will cause agents to behave *as if* they are maximizing an appropriately constructed fitness function (Lehmann et al. 2015). These results however offer little guidance on what (if anything) individuals maximize proximately, i.e., they do not provide specific psychological mechanisms that generate fitness-maximizing behavior. In particular, individuals might get selected to maximize individual fitness, but do this through other-regarding preferences (Akçay et al. 2009). This idea has led to models of preference evolution, where individuals play a given game that has fitness consequences (material payoffs) but where each individual possesses an arbitrary utility function that is genetically determined (Ockenfels 1993; Güth 1995; Akçay et al. 2009; Akçay and Van Cleve 2012; Alger and Weibull 2013). This utility function itself then evolves according to the fitness consequences of the behaviors it generates. Importantly, both the fitness function and the utility function order the outcomes of the social interaction, but these two orderings may be different from each other.

The main result from preference evolution models is that if players can observe each other’s utility functions before choosing an action, then other-regarding preferences may be evolutionarily stable; otherwise, natural selection leads to an utility function that corresponds exactly to the fitness function (Ok and Vega-Redondo 2001; Dekel et al. 2007). This is the same principle that explains the evolution of green-beard genes, where cooperators recognize each other (Robson 1990). A common way for animals to achieve such recognition is repeated interactions where individuals’ behaviors and responses to each other’s behavior is informative of their preferences (Akçay et al. 2009; Akçay and Van Cleve 2012; Jordan et al. 2016). Interactions between relatives also have been shown to promote other-regarding preferences by interacting with such behavioral responses (Akçay and Van Cleve 2012) or recognition of partners (Alger and Weibull 2013).

At the same time, most previous theories that model preference evolution or try to explain cooperation in the laboratory do not take into account that the behavior of humans and other animals is modified by learning based on experienced rewards. Indeed, learning (or initial lack thereof) is usually presented as an alternative to prosocial preferences for explaining behavior in experiments (Binmore 2005). However, as with many social and non-social behaviors consistently produced by a species with a neural system, cooperative behavior must generate positive rewards (in the proximate sense, see below) for an individual (Pearce 2008; Shettleworth 2009; Dugatkin 2010; Schultz 2015). If cooperation is to be observed in those species, then the temporal sequence of cooperation must be consistent with known principles of learning (Sutton and Barto 1998). Moreover, very often in social settings there is uncertainty and variability regarding who is going to be one’s social partner (because the frequency of types changes between generations and because of randomness in the matching process), in which case learning can allow an individual to adapt to its social partners on the timescale of its lifetime. In sum, a theory for the proximate mechanisms of human and animal cooperation is incomplete without accounting for learning at the same time. In a learning context, the question of whether animals have other-regarding preferences thus becomes: can the cooperative act in itself be rewarding?

One may define a reward as an event that generates a particular pattern of activation of neural circuits that induces positive feedback on behavior (Dickinson and Balleine 1994; Pearce 2008; Schultz 2015). Essentially, animals tend to repeat actions that are followed by rewards; this phenomenon constitutes the core of associative learning. Punishments, on the other hand, are stimuli that generate a negative feedback on behavior, whereby actions followed by punishments tend to be avoided in the future. Certain stimuli act as intrinsic rewards (also called primary rewards), which allows an animal to build associations between these intrinsic rewards and new actions or stimuli. Glucose is such an intrinsic reward in many animals: an animal can learn to associate glucose with another stimulus (e.g., a particular fruit), or with an action (e.g., in the laboratory, pulling a lever). Once learning has taken place, the associated stimulus (e.g., the fruit), or the associated action (e.g., pulling a lever) become reward predictors (Niv 2009). Because brain regions involved in decision-making and social cognition project to the mesolimbic reward system (Declerck et al. 2013), it is possible that the part of the brain responsible for social cognition activates this innate reward system. In other words, cooperation may be intrinsically rewarding in the brain. There is evidence that this is true in humans and other primates (Chang et al. 2015). Given the prevalence of cooperation (especially reciprocal cooperation; Taborsky et al. 2016) in many other species, the fact that cooperation is rewarding in the brain is likely to be widespread, although direct neurobiological evidence in other species is scarce. Thus, this basic reward system can be thought of as the proximate basis of learning in social interactions.

From an evolutionary perspective, intrinsic rewards can be viewed as proximate mechanisms that natural selection shapes to make individuals behave in ways that increase fitness. In many cases, intrinsic rewards could be direct proxies of material benefits, as in many models of the evolution of learning (Boyd and Richerson 1988; Josephson 2008; Hamblin and Giraldeau 2009; Arbilly et al. 2010; Katsnelson et al. 2011; Dridi and Lehmann 2014, where the reinforcement term in the equation describing learning is equated to incremental fitness effects). However, in social interactions, intrinsic rewards that are systematically different than one’s own material gains can be evolutionarily stable (Ok and Vega-Redondo, 2001; Dekel et al., 2007; Akçay et al., 2009; Akçay and Van Cleve, 2012; Alger and Weibull, 2013). Natural selection can shape the way social cognition can activate the mesolimbic reward system to take into account stimuli other than one’s own material gain, such as social partners’ payoffs or even abstract social concepts such as fairness and honor, if the resultant behavior is fitness enhancing. Such deviations from a direct mapping from material payoff to intrinsic rewards can evolve either through direct fitness benefits (e.g., because they generate behavioral feedbacks that benefit their carriers; Akçay et al. 2009) or through indirect fitness benefits (Akçay and Van Cleve, 2012). These observations raise the question of how intrinsic rewards that drive learning in social interactions evolve.

In this article, we present a model of the evolution of such intrinsic rewards when individuals interact in the Prisoner’s Dilemma game, where they have the choice between cooperation and defection. To capture general learning processes in humans and other animals, we model learning as a basic trial-and-error process where individuals repeat actions followed by rewards and avoid actions followed by punishments. In our model, individuals interact in games whose material payoffs determine fitness. Instead of learning according to the real material payoffs, an individual associates to each game outcome a genetically determined utility, which is used as the intrinsic reward/punishment for that particular outcome. For example, other-regarding individuals may associate positive utilities to outcomes where their partner obtains a positive material payoff, and thus might learn to cooperate as an intrinsically rewarding action. This decoupling of material payoffs and rewards allows us to address the question of how rewards evolve in social interactions. We look for the evolutionarily stable utility functions when individuals interact repeatedly in a game whose material one-shot payoffs determine the 2-person Prisoner’s Dilemma game.

## Model

### Social interactions and rewards

We consider an evolutionary model of repeated pairwise games in a large, well-mixed population of learners with non-overlapping generations. Every generation of the evolutionary process consists of a sequence of interaction rounds, *t* = 1, 2, …, *T*. At each generation just before *t* = 1, individuals in the population are randomly matched in pairs, and each pair remains together for the entire duration of the game (until *t* = *T*). Hence, individuals are playing a repeated game with their partner. The one-shot game, played at every time *t*, is a Prisoner’s Dilemma game with two possible actions, cooperate, *C* (or action 1), or defect, *D* (or action 2). The one-shot material payoffs for individual *i* are denoted *π*_*i*_(*C, C*) = *b* − *c, π*_*i*_(*C, D*) = −*c, π*_*i*_(*D, C*) = *b, π*_*i*_(*D, D*) = 0, where the first element in parentheses of *π*_*i*_(*a*_*i*_, *a*_−*i*_) denotes player *i*’s action (*a*_*i*_), and the second element denotes his opponent’s action (*a*_−*i*_). We assume also that *b* > *c* > 0. The sequence of material payoffs ultimately determines fitness (see below for details on how fitness is evaluated).

At every interaction round *t*, each individual in every pair chooses an action. Individual *i*’s action at time *t* is denoted *a*_*i,t*_ and his opponent’s action is *a*_−*i,t*_. After actions are chosen, both players observe the outcome (*a*_*i,t*_, *a*_*−i,t*_) and subjectively evaluate how good the outcome was, which is genetically determined. We call this subjective evaluation the utility function of a player, which may be different than the actual material payoff, *π*_*i*_(*a*_*i,t*_, *a*_−*i,t*_), obtained at time *t*. This utility (rather than the material payoff) determines the reward sensation of a game outcome, and this reward is used by an individual to learn his strategy in the repeated game (see below for details about the learning process). Specifically, the genotype of each individual *i* associates to each outcome (*a*_*i*_, *a*_−*i*_) an utility *u*_*i*_(*a*_*i*_, *a*_−*i*_)that can take any negative or positive real value. We say that the utility is a reward if it is positive *u*_*i*_(*a*_*i*_, *a*_−*i*_) > 0, while we call it a punishment if *u*_*i*_(*a*_*i*_, *a*_−*i*_) < 0. Hence a genotype consists of the four utilities *u*_*i*_(*C, C*), *u*_*i*_(*C, D*), *u*_*i*_(*D, C*), *u*_*i*_(*D, D*). We can arrange these four utilities in a matrix according to the game outcomes, which we call the utility matrix of individual *i* (Fig. 2A). However, evolutionarily speaking, it is easier to think of these utilities as the vector **u**_*i*_ = (*u*_*i*_(*C, C*), *u*_*i*_(*C, D*), *u*_*i*_(*D, C*), *u*_*i*_(*D, D*)); below we also use the more compact notation **u** = (*u*_11_, *u*_12_, *u*_21_, *u*_22_), dropping the player’s index. The state space is thus ℝ^4^. Our interest in this paper is to find the evolutionarily stable utility vector **u**\*. To do so, we need to know the fitness *f*(**u**_*i*_) of an individual with utility **u**_*i*_. To arrive there, we first need to specify how the utility vectors of a pair of players determine behavior in the repeated game.

**Figure 1:**
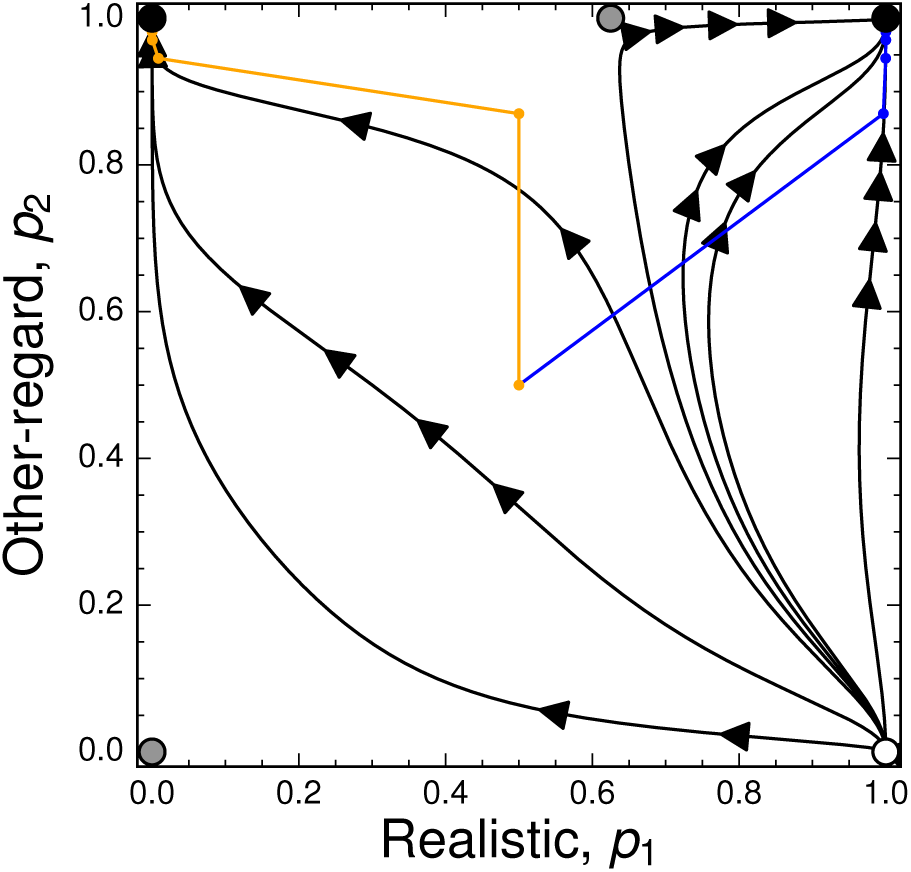
Solution trajectories (black) and stochastic trajectories (colored lines) for the behavioral interaction between Realistic and Other-regard. On the *x*-axis is represented the probability that Realistic cooperates (*p*_1_), while on the *y*-axis, this is the probability that Other-regard cooperates (*p*_2_). The stochastic trajectories are started from the center of the state space 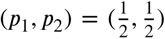 and dots on it represent interaction rounds between the players. Circles represent equilibria: a white-filled circle is a source (both associated eigenvalues are positive); a gray-filled circle is a saddle (one positive and one negative associated eigenvalue); a black circle is a sink (both associated eigenvalues are negative).

**Figure 2:**
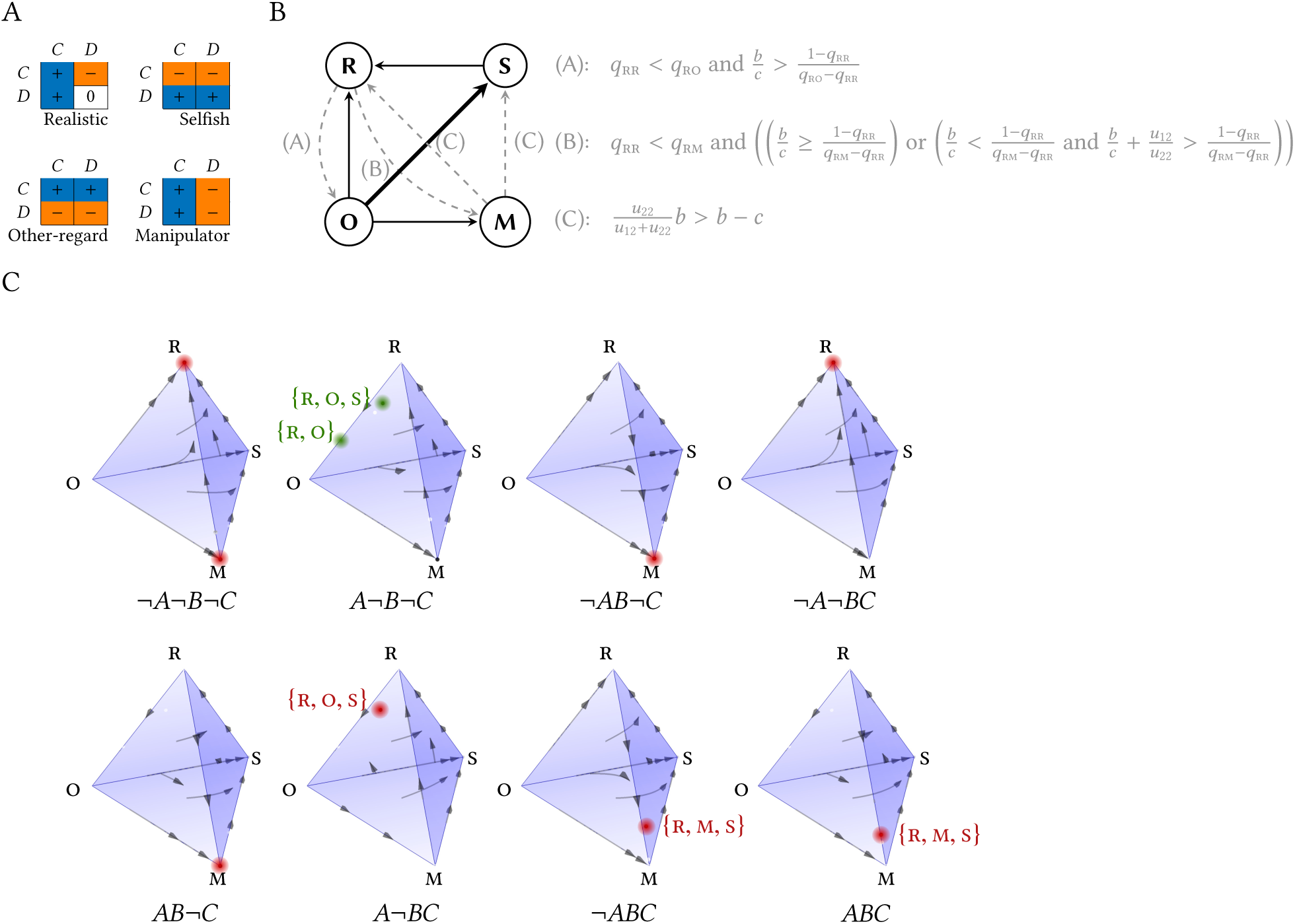
Replicator dynamics for the competition between Realistic, Other-regard, Manipulator, and Selfish. (A) The 4 strategies considered in the analytical model. A strategy is defined by the outcomes to which it associates a positive or negative utility. The first row/column corresponds to Cooperate and the second row/column to Defect. Utilities are to row-player. (B) Pairwise invasion diagram between the four strategies Realistic, Other-regard, Manipulator, and Selfish, and associated invasion conditions. A plain directed edge from node X to node Y means that strategy Y always invades a monomorphic population of X (but does not necessarily reach fixation). A dashed directed edge from node X to node Y means that Y can invade X under certain conditions (A, B, and C) on the model parameters. When a given strategy can be invaded by more than one other strategy, a thick edge designates the best response. Note that all combinations of these three conditions are possible. (C) Classification of phase portraits for the replicator dynamics in the 4-strategy game defined by the competition between Realistic, Other-regard, Manipulator, and Selfish. Each subfigure is a drawing of the 4-simplex. At each vertex, one of the four strategies is at frequency 1: Realistic at the top, Manipulator at the bottom-front, Other-regard at the back left, and Selfish at the back right. The letters *A, B*, and *C* refer to the conditions in Panel B, where the symbol “¬” denotes logical negation. For instance, the subfigure labeled ¬*ABC* is drawn for parameter values such that condition *A* is not true, but conditions *B* and *C* are true. Red dots denote locally-stable equilibria, i.e. possible outcomes of natural selection. To disambiguate the 3D view, red labels in curly braces indicate the set of strategies present at an equilibrium. In the subfigure for the case *A*¬*B*¬*C*, the green dots are two alternative outcomes: {R, O} occurs when Selfish cannot invade this polymorphism; if Selfish does invade this polymorphism, {R, O} beaomes unstable and {R, O, S} stable. The condition for this to happen is given by eq. A11 in Appendix A.

### Learning

We assume that individuals learn to play the game according to a simple trial-and-error procedure. We use a standard model of learning dynamics (Sutton and Barto 1998; Dridi and Lehmann 2014), except that actions are reinforced according to the subjective utilities of a game outcome *u*_*i*_(·), rather than the objective material payoff *π*_*i*_(·). At every time *t*, an individual *i* holds in memory action values *V*_*i,t*_(*a*_*i*_) for both actions *a*_*i*_ ∈ {*C, D*} and chooses to cooperate at time *t* with a probability, *p*_*i,t*_, that depends on its action values {*V*_*i,t*_(*C*), *V*_*i,t*_(*D*)}. Adapting an existing model of the evolution of learning rules (Dridi and Lehmann 2015), the learning rule of individual *i* in our model is to update action values according to

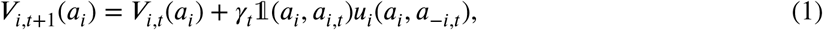
where 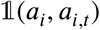 is an indicator variable that equals 1 if *a*_*i*_ = *a*_*i,t*_, and 0 otherwise, and *γ*_*t*_ ∈ (0,1) is a dynamic learning rate. This learning rate is decreasing as the game proceeds, which implies that the initial rounds of interaction are critical in determining the stable outcome of the learning process. Such a condition ensures that learning converges during an individual’s lifetime (Benaïm, 1999; Dridi and Lehmann, 2014) and can be justified by the fact that the game faced by the individuals is constant. Finally *u*_*i*_(*a*_*i*_, *a*_−*i,t*_) is the utility to *i* if he plays *a*_*i*_ given that his opponent plays *a*_−*i,t*_ at time *t*.

We then assume that individuals want to choose the action with highest value *V*_*i,t*_(*a*_*i*_), but also have some tendency to explore the action with smaller value. A widely used choice rule to capture this principle is the logit-choice function,

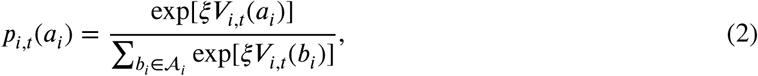
where *ξ* > 0 is the exploration parameter (the inverse 1/*ξ* can be seen as the noise level if we interpret this model as perturbed maximization of action values, Hofbauer and Sandholm, 2002) in choosing actions. In our case, there are two actions, *C* and *D*, hence eq. 2 is a sigmoid function, which can be thought as a generalization of the threshold rule that chooses the action with greater value *V*_*i,t*_(*a*_*i*_). Eq. 2 approaches such threshold function when *ξ* gets larger.

The behavioral interaction between a reinforcement learner with utilities **u** = (*u*_11_, *u*_12_, *u*_21_, *u*_22_) and another reinforcement learner with utilities **v** = (*υ*_11_,*υ*_12_, *υ*_21_, *υ*_22_) is what we need to analyze in order to compute fitness. By a slight abuse of notation, we denote these two players *u* and *υ* and their probabilities to cooperate by *p*_*u*_ and *p*_*υ*_ respectively. Stochastic approximation theory can be used (see e.g., Dridi and Lehmann 2014) to show that the long-run learning dynamics (eqs. 1–2) for a pair of learners can be described as

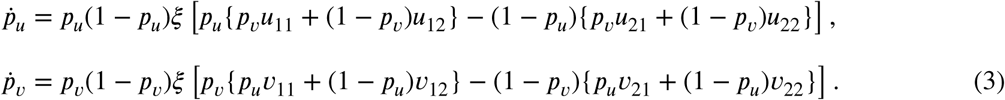

Eq. 3 displays ten generic behavioral equilibria (Fig. B1 in Appendix B). Depending on the specific values of **u** and **v**, one or more of these equilibria may exist. Note that because the original dynamic is stochastic, when the corresponding deterministic system admits several locally stable equilibria, the stochastic dynamics may reach any of these equilibria. It turns out that the theory of stochastic approximations is almost silent about which particular equilibrium will be reached. These lock-in probabilities will however play an important role for the evolutionary stability of the different utility functions we will study below.

Another important fact about the behavioral dynamics is that the stability of the possible behavioral equilibria is very much dependent on the signs of utilities of the individuals involved in an interaction. In particular, one has that a pure equilibrium is locally stable if and only if both players have a positive utility (making it a *reward*) for this outcome. The implication of this is that if at least one player has a negative utility for the outcome, then this outcome is unstable. In other words, if players *u* and *υ* do not “agree” on preferred outcomes, then a pure behavioral equilibrium cannot be stable. This intuitive result is mathematically true because the eigenvalues of the Jacobian matrix associated to eq. 3 evaluated at a pure outcome (*i, j*) are simply

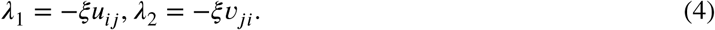

This fact has important evolutionary consequences, as will be detailed below when we analyze interactions between individuals with particular utility functions. In particular, it allows us to classify different utility functions by their sign for each of the four outcomes.

### Fecundity

Assuming that interactions last long enough (*T* → ∞), we define the fecundity of individual *i* as being proportional to the average material payoff obtained at equilibrium of the learning process, i.e.

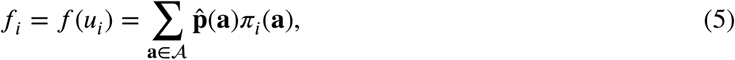
where 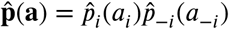 is the equilibrium probability of outcome **a** = (*a*_*i*_, *a*_−*i*_). The sum in eq. 5 is taken over the set of possible game outcomes, 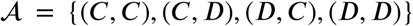. We call 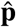 the behavioral equilibrium. Importantly, while the utility function does not appear on the right-hand side of eq. 5, we still defined it as *f*(*u*_*i*_) because the equilibrium choice probabilities of a player, 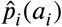, implicitly depend on the utility function of player *i*, as will become clearer when we derive expressions for the behavioral equilibria below.

The fecundity *f*(**u**) depends on the outcome of the learning dynamics, and is therefore not continuous in **u**, which renders difficult a full analytic treatment of the model. To overcome this problem we adopt two complementary approaches. First, we focus on a smaller number of utility functions that are relevant to our original question of the evolution of other-regarding preferences. Second we run evolutionary simulations of the full model to have a more comprehensive view of our model.

## Results

### Analytical results for 4-strategy competition

We first consider the evolutionary dynamics of four possible utility functions that are represented in Fig. 2A using the replicator dynamics (for details of the analysis, see Appendix A):

- The **Realistic** function, which associates to outcomes an utility of the same sign than the real material payoff. This type of utility function is the “default” utility function, used in virtually all models of the evolution of learning (Boyd and Richerson 1988; Josephson 2008; Hamblin and Giraldeau 2009; Arbilly et al. 2010; Katsnelson et al. 2011; Dridi and Lehmann 2014). It takes as a special case the material payoff function, i.e., *u*_*i*_ = *π*_*i*_. It is the function that evolves when interactions between players are completely anonymous, one-shot, and there is no assortment in the matching process (Ok and Vega-Redondo 2001; Dekel et al. 2007).
- The **Other-regard** function, which associates positive utility to the outcomes where the opponent obtains a strictly positive payoff. In other words, this strategy associates positive utilities only to the outcomes where it cooperates.
- The **Selfish** function, which associates positive utility to the outcomes where it defects.
- The **Manipulator** function, which associates positive utility only to the outcomes where its opponent cooperate. The name of this utility function stems from the fact that it will drive a compliant opponent (who associates positive utility to all outcomes) to cooperate.

We first construct the fitness matrix for the evolutionary game in Table 1 by considering the stable equilibria of learning dynamics for all possible pairwise matchings between the four strategies (described in detail in the Appendix A; see also Fig. 1 and Fig. B2). For the four strategies we consider in this section, no more than two behavioral equilibria are locally stable at the same time. It turns out that in all cases where two equilibria are locally stable, one of them is mutual cooperation, (1, 1) (Fig. 1 and Fig. B2). Because the underlying learning model is stochastic (eqs. 1–2), the lock-in probability in the cooperative equilibrium (1, 1) will affect the fitness and hence the evolutionary competition between the four strategies we are considering. However, there is no general technique to obtain an expression of the lock-in probability. At this point, we leave these probabilities unspecified, and denote by *q*_*uυ*_ the probability that an interaction between strategy *u* and strategy *υ* leads to the cooperative equilibrium (1, 1). For instance, two Realistic individuals can learn mutual cooperation, (1, 1), or mutual defection, (0, 0). The probability that an interaction between two Realistic individuals leads to mutual cooperation is thus denoted *q*_RR_; the probability of locking in the defective equilibrium is 1 − *q*_RR_.

### Evolutionary dynamics for the Prisoner’s Dilemma

We use the replicator dynamics (Taylor and Jonker 1978, eq. A6 in Appendix A) to describe the competition between Realistic, Other-regard, Manipulator, and Selfish, with the evolutionary game given in Table 1. Determining the outcome of the replicator dynamics is dependent on the parameters *b* (benefit to a receiver of a cooperative act), *c* (cost of cooperating), and the lock-in probabilities in the cooperative equilibrium *q*_RR_, *q*_RO_, *q*_RM_, *q*_OM_, for the different behavioral interactions where several equilibria are locally stable.

**Table 1:** Evolutionary fitness matrix amongst the 4 strategies considered in the analytical model.

|  | R | O | M | S |
| --- | --- | --- | --- | --- |
| R | $q_{RR}(b - c)$ | $q_{RO}(b - c) + (1 - q_{RO})b$ | $q_{RM}(b - c) + (1 - q_{RM})\left(b\left(\frac{u_{22}^M}{u_{12}^M + u_{22}^M}\right)\right)$ | 0 |
| O | $q_{RO}(b - c) + (1 - q_{RO})(-c)$ | $b - c$ | $q_{OM}(b - c) + (1 - q_{OM})(-c)$ | $-c$ |
| M | $q_{RM}(b - c) + (1 - q_{RM})\left((-c)\left(\frac{u_{22}^M}{u_{12}^M + u_{22}^M}\right)\right)$ | $q_{OM}(b - c) + (1 - q_{OM})b$ | $b - c$ | $-c\frac{u_{22}^M}{u_{12}^M + u_{22}^M}$ |
| S | 0 | $b$ | $b\frac{u_{22}^M}{u_{12}^M + u_{22}^M}$ | 0 |

Note: $q_{ij}$ denotes the probability that a behavioral interaction between strategy $i$ and strategy $j$ leads to mutual cooperation at a behavioral equilibrium. The expression $u_{ab}^i$ denotes the utility of strategy $i$ for the game outcome where it chooses action $a$ and its opponent chooses $b$ .

We first find that, although (Selfish, Selfish) is always a weak Nash Equilibrium (NE) of the evolutionary game between the four strategies, regardless of parameter values, it is never evolutionary stable (Fig. 2 and Table 1). This is because Selfish gets invaded by Realistic, which learns to defect against Selfish, but cooperates with itself. On the other hand, the strategy Other-regard is also always invaded by every other strategy in pairwise competitions, although it can be part of a mixed equilibrium, as we will see below. All other important results depend on the parameters of the model, and three basic conditions on the parameters help classify the possible evolutionary outcomes (conditions A, B, and C in Fig. 2B). For certain parameter values, Realistic can be an evolutionarily stable strategy when the benefit-to-cost ratio is sufficiently low (conditions A and B in Fig. 2B). Also, for other parameter values, Manipulator can be evolutionary stable (condition C in Fig. 2B). Note that these conditions are not mutually exclusive, so both Realistic and Manipulator can be evolutionarily stable at the same time (Fig. 2C). When at least one of Realistic or Manipulator is not evolutionarily stable, then we obtain polymorphic equilibria. In such polymorphic equilibria, we have either three strategies (there is an equilibrium with Realistic, Other-regard, Selfish, and an equilibrium with Realistic, Manipulator, and Selfish) or two strategies (Realistic and Other-regard), the common feature of these being that Realistic is always present (Fig. 2C). We note here that Other-regard can only be present when Realistic is present. Moreover, according to condition A in Fig. 2B, Realistic should cooperate more often with Other-regard than with itself for the latter to make part of an equilibrium.

## Simulations

### Freely evolving utilities

The above analysis pertains to “internal stability” (cf. Eshel 1996) among a restricted class of utilities in the short-term. However, utilities in our model comprise a 4-dimensional vector of continuous evolutionary strategies. Therefore, it is natural to ask how populations, impelled by natural selection, move through such a strategy space in the long-term. To do that, we conducted stochastic evolutionary simulations, in which we introduce new mutations from a much less constrained strategy space to a monomorphic population and use results from well-established population genetic theory (Van Cleve, 2015) to calculate the invasion success of this mutant. This allows us to explore the strategy space in a computationally efficient manner.

In our simulations, the lock-in probabilities in behavioral equilibria, which played a critical role in determining the evolutionary outcome in the above 4-strategy model, will no longer be parameters but will have a value that depends on the utilities of the particular strategies involved in behavioral interactions. Our evolutionary simulations consist of the trait substitution sequence of adaptive dynamics. Namely, we assume that the genotype of an individual, **u** = (*u*_11_, *u*_12_, *u*_21_, *u*_22_), is supported by one locus, and that the population is always monomorphic. At each iteration, we propose a mutation and determine whether the mutant invades the resident population using eqs. 41–42 of Van Cleve (2015), which is calculated for Wright’s island model (in our case, the population is panmictic, or there is only one deme). We performed our evolutionary simulations for various values of the benefit-to-cost ratio, *b*/*c* as well as different values of game duration, *T*.

To describe the results and in order to represent the four utilities at the same time, we classified all strategies according to the sign of their utilities (as we demonstrated above, these signs provide necessary conditions on the possible behavioral equilibria), which results in 2^4^ = 16 classes of strategies (because each of the four utilities has two possible signs). We can first look at the proportion of time a simulation run spends in each of the 16 strategy classes, which is an approximation of the stationary distribution of the evolutionary dynamics. We find that 6 strategies are consistently represented more than 10% of the time in the stationary distribution: Selfish, Avoid Sucker’s Payoff, Manipulator, Matcher, Pareto, Anti-Cooperation. Avoid Sucker’s Payoff (AS) is similar to Realistic except that it has a positive utility for mutual defection instead of a 0; AS produces the same behavioral equilibria as Realistic when paired with other strategies (Fig. B3). Matcher has positive utilities only for outcomes where its own action matches that of its opponent, thus the pure outcomes it can learn are mutual cooperation or mutual defection. Pareto has positive utility only for the outcome of mutual cooperation, thus it will never learn full defection and will learn mutual cooperation against any opponent who is also willing to do so. Finally, Anti-Cooperation is the exact opposite of Pareto, as it has positive utilities for all outcomes except for mutual cooperation; this utility matrix cannot learn mutual cooperation, generally learns to defect, but may be exploited by exploiting strategies such as Manipulator (Fig. B3). In Fig. 3, we show results for various benefit-to-cost ratios, *b*/*c*, which leads to two main observations.

**Figure 3:**
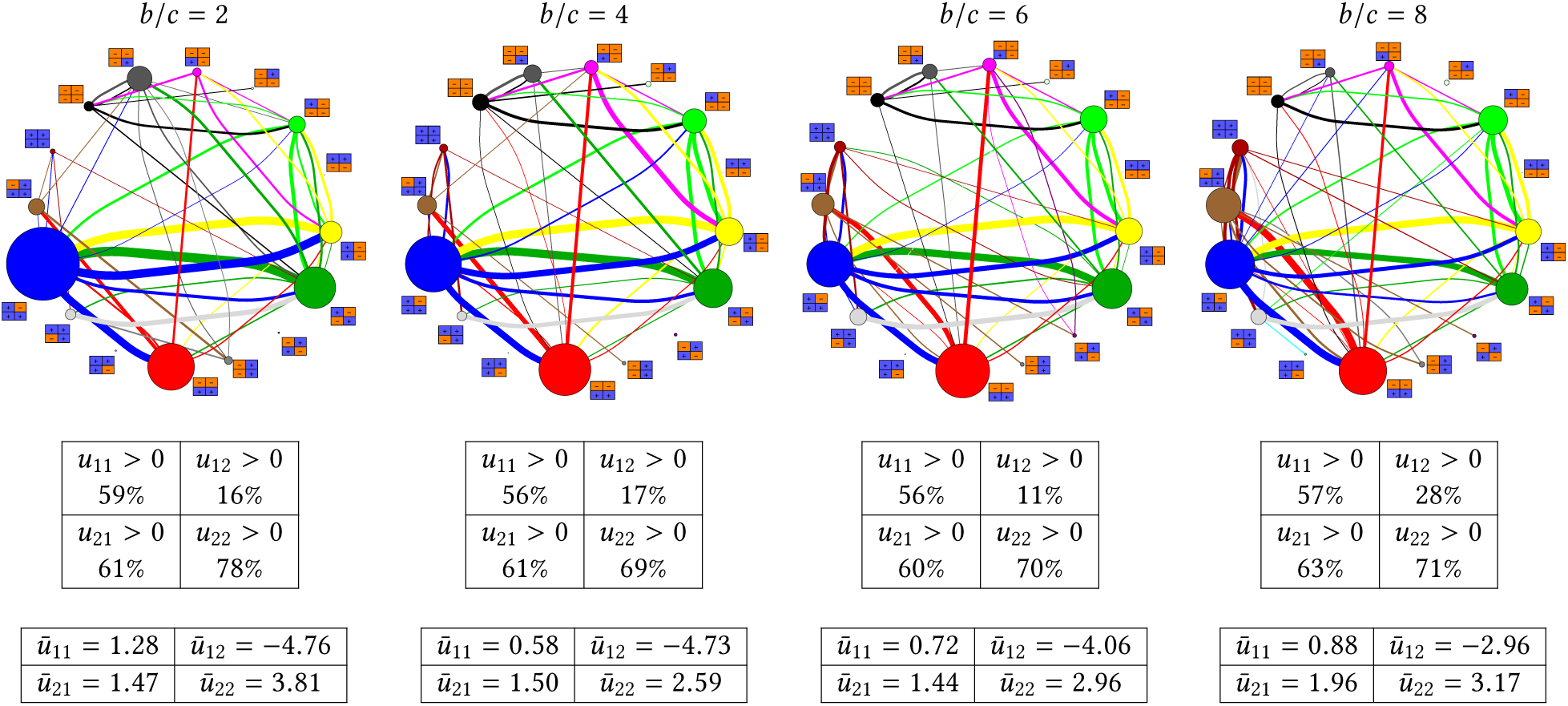
Invasion analysis and utilities in simulations for various benefit-to-cost ratios (*b*/*c*). The top row shows the invasion graph between the 16 classes of strategies defined by their utilities’ sign (see main text), the second row shows the proportion of time each utility was positive in a simulation run, the third row shows the time average of utilities. In the invasion graph, near each node we show the utility matrix of the corresponding strategy, with blue cells indicating a positive utility, and orange cells indicating a negative utility. The utility matrix is oriented as in Fig. 2. The size of the nodes is proportional to the amount of time a simulation run spends in the corresponding strategy class. The edges are colored according to the invader strategy and thus indicate the direction of the edges. Edge thickness is proportional to the number of invasions that occured between a pair of strategies (and we do not show edges between pairs of strategies for which the number of invasions was less than 10). Parameter values: min *u* = −10; max *u* = 10; *ξ* = 2; *c* =1; *T* =150; Population size *N* = 2000.

The first observation is that our simulations confirm the overall pattern in the analysis of the replicator dynamics, where at low values of *b*/*c* the strategy AS (corresponding to the Realistic strategy in the analytical model) experiences few invasions. As *b*/*c* increases, more strategies are able to invade AS, and consequently the frequency of AS declines (Fig. 3 and Fig. 4). In particular, if we analyze the invasions between the 6 dominant strategies in our simulations (Fig. B4), we find that Manipulator, Matcher, and Pareto invade AS only for sufficiently high *b*/*c*. All these strategies have a positive utility for mutual cooperation; they also have a negative utility for the sucker’s outcome (*u*_12_ < 0). The success of AS and of cooperative strategies more generally yields an average utility matrix of the AS type (Fig. 3), where average utilities are ordered as *ū*(*D, D*) > *ū*(*D, C*) > *ū*(*C, C*) > *ū*(*C, D*), which is different than the ordering of the material payoffs, *π*(*D, C*) > *π*(*C, C*) > *π*(*D, D*) > *π*(*C, D*). The strategy Pareto increases in frequency in the stationary distribution for increasing *b*/*c* (Fig. 4A), as the analysis shows that it invades AS for high enough *b*/*c* (Fig. B4). Strategies that are able to invade AS (Manipulator, Matcher, Pareto) can mutually invade one another and we indeed observe that an important number of invasions occur between AS, Manipulator, Matcher, Pareto (Fig. 3). As a consequence of the increasing success of strategies that positively value cooperation as a function of *b/c*, we observe that the overall cooperation frequency in the population increases for increasing *b*/*c* (Fig. 4B). Even though previous work has shown that cooperative strategies in the iterated Prisoner’s Dilemma can be evolutionary robust (Stewart and Plotkin 2013), we could not expect this for the particular type of learning strategies that we have decided to study.

**Figure 4:**
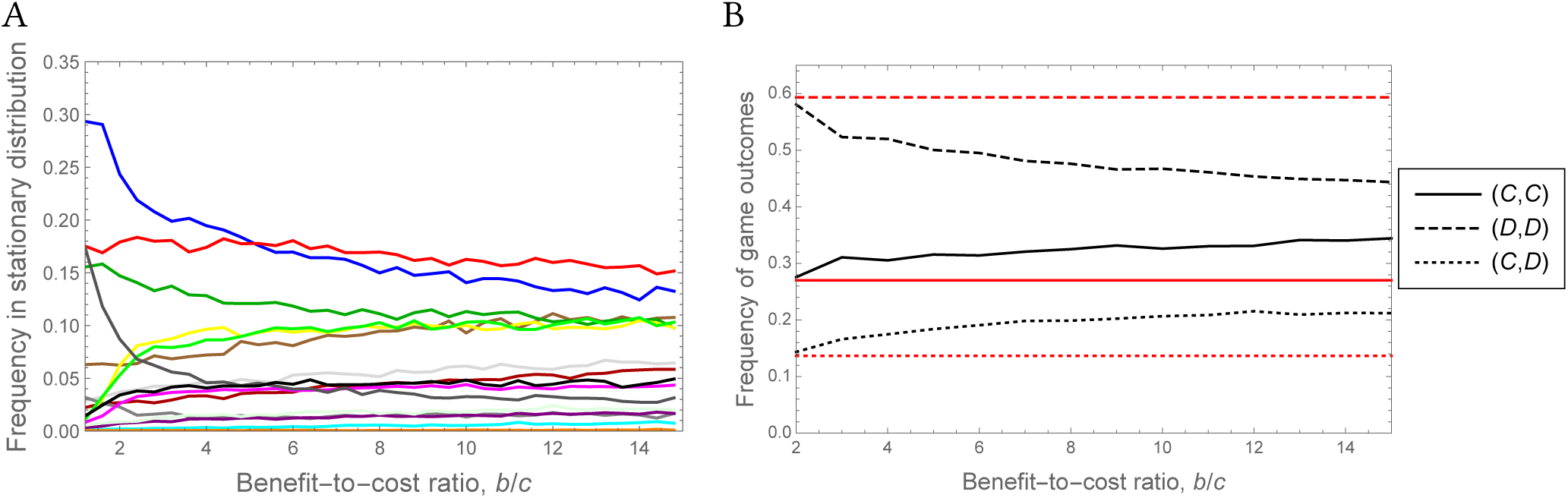
Effect of the benefit-to-cost ratio, *b*/*c* on the stationary distribution of strategies and effective cooperation. (A) Proportion of time the simulation spends in each of the 16 strategy classes as a function of *b*/*c*, which is a measure of the stationary distribution. Colors of strategies are as in Fig. 3 (see also Fig. B5). (B) Time average of the frequency of game outcomes in a simulation run as a function of *b*/*c* (black lines; note that (*C, D*) and (*D, C*) are the same outcome). The red lines show the expected frequency of the corresponding game outcomes in the population if we draw strategies randomly from a uniform distribution. Parameter values are as in Fig. 3.

Contrasting the apparent success of conditionally cooperative strategies, a second major feature of our simulations is the success of Selfish. For all *b/c*, the simulation spends approximately 15-20% of the time in this strategy class, and for high *b*/*c* this makes Selfish the most represented strategy class in the stationary distribution (because of the decline of AS; Fig. 3 and Fig. 4A). Although this result could not be anticipated from our analysis of the replicator dynamics above, it is still consistent with the fact that Selfish was relatively stable (only Realistic could invade it). Analytically considering the invasion conditions between the 6 dominant strategy classes in the simulations (Fig. B4) reveals that Selfish is also relatively stable in this set, with only AS and Matcher being able to invade it. In our simulations, AS invades more frequently Selfish than Matcher does because, with our mutation scheme, an AS mutant is much more likely than a Matcher mutant to occur in a Selfish population given that we draw mutations from a doubly exponential distribution centered at the resident phenotype.

A final observation regarding strategy classes is that Anti-Cooperation is relatively successful for high *b*/*c* (Fig. 4A). Our invasion analysis in Fig. B4 shows that this strategy, despite having a positive utility for the sucker’s outcome, compensates by exploiting certain cooperative strategies, such as Pareto. At low *b*/*c*, Anti-Cooperation gets exploited by strategies that have positive utilities for defection (such as AS or Selfish; Fig. B3) but as *b*/*c* increases, Anti-Cooperation becomes more stable against these strategies, which explains why it makes part of an important proportion of the stationary distribution of the evolutionary dynamics.

In order to verify whether these results were sensitive to the length of the repeated game, we ran additional simulations for lower values of *T*. Our analytical results were obtained under the assumption that *T* is large enough so that learning reaches an equilibrium during individuals’ lifetime, and our standard simulations were run for *T* = 150. When using *T* = 50, we essentially obtain the same results in terms of the stationary distribution of strategies (Fig. B9A,B). We needed to decrease the duration of the game to *T* = 10 to obtain different results (Fig. B9C,D). Namely, in this case we find that Selfish is the mode of the stationary distribution for all benefit-to-cost ratios. Otherwise we observe a similar pattern than for higher *T* values, with AS being represented more than other strategies for low *b*/*c*, but slowly decreasing as *b*/*c* increases (Fig. B9C). For this low *T* value, we also observe that the strategies Manipulator, Matcher, Pareto and Anti-cooperation, which previously grew in frequency for increasing *b*/*c* still do so. The apparent success of Selfish for low *T* is due to the fact that for this duration of the game, learning cannot reach an equilibrium for all strategies, and strategies that can learn multiple stable behavioral outcomes may wander between equilibria. In contrast, Selfish can only learn defection, irrespective of the opponent and its convergence to the equilibrium occurs faster. Hence the strategies that previously (i.e. for *T* = 150 or *T* = 50) succeeded in both cooperating with cooperators but defecting with Selfish now fail to learn fast defection against Selfish. In the Appendix B we illustrate this phenomenon for interactions between the AS strategy and Selfish (Fig. B10).

### Utilities that explicitly depend on material payoffs

In this section we perform additional simulations by constraining the utility function to be dependent on the material payoffs of the focal player and its opponent. This allows us to address more directly the question o^f^ whether (and, if any, what type of) other-regarding preferences evolve in our model. Specifically, for any game outcome **a** = (*a*_*i*_, *a*_−*i*_), we consider utility functions of the form

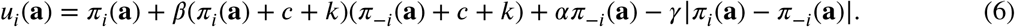
where (*α, β, γ*) are player *i*’s genetically determined parameters. In this section we will be interested in the evolution of these three parameters. In eq. 6, *c* is the negative of the sucker’s payoff (−*c*) and is added to the realized payoff to ensure that the term multiplied by *β* is always positive. The parameter *k* is here to allow the utility to vary as a function of *β*. Our utility function then measures the extent to which an individual is “additively” other-regarding (*α* ∈ [−1, 1]), the extent to which he is “multiplicatively” other-regarding (*β* ∈ [−1, 1]), and inequity aversion (*γ* ∈ [−1, 1]), Even though this utility function can realize all of the 16 possible utility matrices discussed above, the structure of the phenotype space changes as compared to the above simulations where we let the utility matrix evolve in an unconstrained way (Fig. B5).

Our simulations with the utility function in eq. 6 show that the selection pressure on other-regarding preferences increases with *b*/*c* (Fig. 5A and Fig. B8). The average value of *β* is close to 0 for low enough *b*/*c*, but suddenly increases at a threshold value of *b*/*c*. For these higher *b*/*c* values, the average *β* is approximately 0.5, indicating the evolution of multiplicative other-regard. The average values of *α* and *γ* are negative for low *b*/*c*, indicating respectively a combination of competitive preferences (valuing negatively other’s success) and inequity aversion. Both *α* and *γ* decrease in magnitude as *b*/*c* increases, but remain negative. This is a consequence of the fact that the selection pressure on *α* and *γ* decreases with increasing *b*/*c*, because the absolute difference between the temptation to defect, *b* and the sucker’s payoff, −*c*, decreases. This pattern is accompanied by a general increase in the utility for mutual cooperation as a function of *b*/*c* (Fig. B7A in the SI). For high *b*/*c*, mutual cooperation becomes the preferred outcome of the evolutionarily stable utility function and mutual defection the least preferred outcome. In agreement with the above simulations for freely evolving utilities, AS is the dominant utility matrix for low *b*/*c*. The “Compliant” utility matrix (with all four utilities positive) becomes the dominant one for high *b*/*c* (Fig. B7B in the SI).

**Figure 5:**
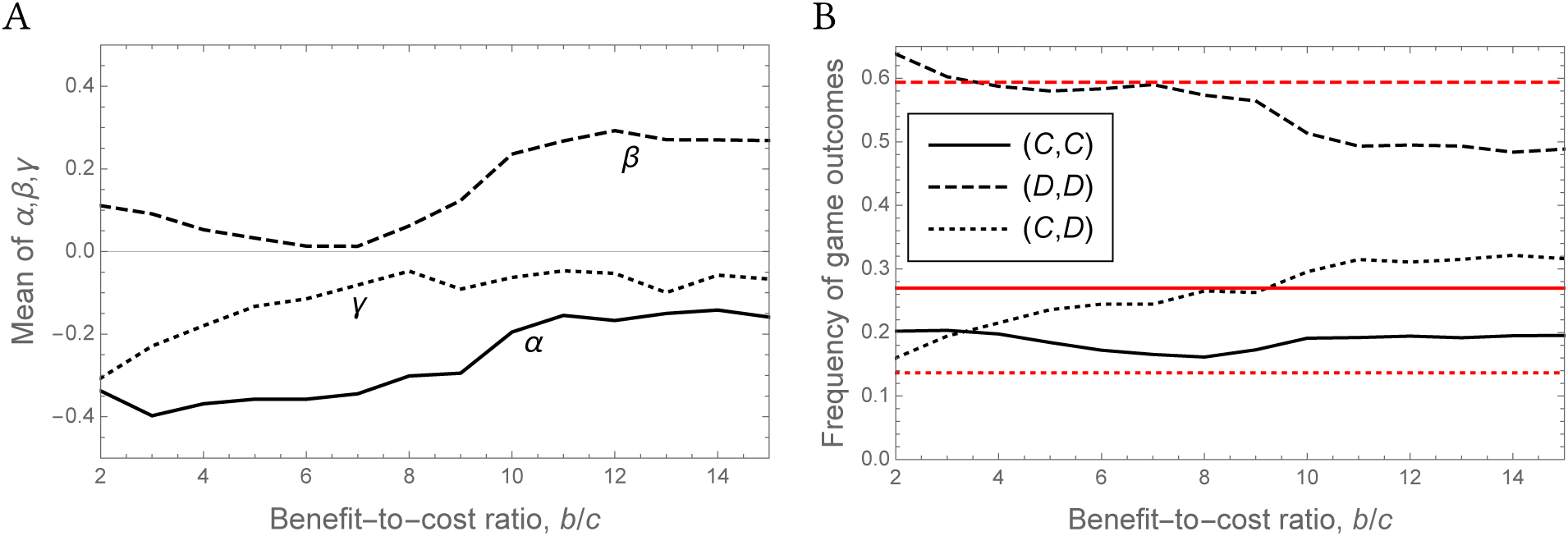
Results for the model where the utility depends explictly on material payoffs (eq. 6 with *k* = 2) as a function of the benefit-to-cost ratio, *b*/*c*. (A) Time average of *α, β, γ* in a simulation run. (B) Time average of the frequency of game outcomes in a simulation run (similar to Fig. 4).

Even though other-regarding preferences evolve for sufficiently high *b*/*c*, this is not accompanied by the evolution of increased effective mutual cooperation, even though the frequency of mutual defection decreases. This decrease in mutual defection is due to an increase in the asymmetric (*C, D*) outcome (Fig. 5B). Overall cooperation is thus increasing but individuals do not coordinate on cooperating at the same time. This can be explained by the fact that the Compliant utility matrix that evolves for high *b*/*c* can learn any outcome (all pure equilibria are stable when all utilities are positive). However, the fact that mutual defection is the least preferred outcome implies that the probability to learn this equilibrium will be the lowest of the four outcomes.

## Discussion

### Evolution of rewards for prosocial learning

We presented a model of how intrinsic rewards that drive learning in social interactions evolve. Rewards capture the intrinsic preferences of individuals over states of the world, and constitute the fundamental building block of reinforcement learning. Because all behaviors are in part influenced by learning, modeling the evolution of social behaviors in animals requires that we take into account how behavior is generated through learning within an individual’s lifespan. Within this framework, we developed *a* model to provide insight into the question of whether other-regarding preferences support the evolution of cooperation, under the constraint of reinforcement learning. While previous theoretical work tended to either ignore or oversimplify learning mechanisms (Ok and Vega-Redondo 2001; Dekel et al. 2007; Akçay et al. 2009; Akçay and Van Cleve 2012; Alger and Weibull 2013), our model tries to account for the increasing empirical evidence that reward processing and learning are critical aspects of prosocial preferences (Fehr and Camerer 2007; Declerck et al. 2013; Ruff and Fehr 2014). Overall, our results indicate that multiple preference functions can be evolutionarily stable when individuals interact repeatedly in the Prisoner’s Dilemma. In particular, we find that evolutionarily successful preferences are of two general types: (1) those that have a positive utility for mutual cooperation but a negative utility for being exploited; (2) selfish preferences that associate positive utilities to outcomes where their carriers defect, and have negative utility for cooperation. This is true in both our analytical results in replicator dynamics and the numerical simulations in the whole strategy set. Further simulations show that other-regarding preferences evolve for sufficiently high benefit-to-cost ratio.

A majority of the empirical evidence for the existence of other-regarding preferences come from experiments performed by economists with humans participants. Economic theory relies on the concept of utility to capture behavior, but the utility function of an individual is by definition an internal construct that is difficult to access (Fehr and Camerer 2007). In the context of learning, utility can be equated to reward, because rewards are at the core of repeated behaviors (Schultz 2015). Empirically, one way to try to access the utility or reward function is to observe the pattern of activation in the brain when individuals make decisions. One of our main findings is that positive preferences for cooperation are evolutionarily prevalent. This finding is interesting when paralleled with neuro-behavioral studies of social decision-making that reveal that cooperation can generate rewards in the human brain, which seems consistent with the positive utility of winning strategies for mutual cooperation found in our model (Fehr and Camerer 2007; Declerck et al. 2013; Ruff and Fehr 2014). Additional empirical and theoretical work focusing on the cooperative behavior, but not on the preferences generating it, has already been conducted based on the premise that individuals may act prosocially in order for others to recognize their willingness to cooperate (Gintis et al. 2001; Jordan et al. 2016) or that cooperation could rely on fast decision-making implying a possible intrinsic preference for cooperation (Rand et al. 2012; Tinghög et al. 2013). Although these studies do not directly measure or model directly social preferences, they are in principle compatible with evolved preferences that find cooperation intrinsically rewarding. More generally, our results suggest a possible psychological mechanism for reciprocal cooperation in other animal species (Taborsky et al. 2016). Indeed the evolutionarily stable utility functions in our model that positively value mutual cooperation produce behavioral dynamics that resemble the dynamics of reciprocal strategies such as tit-for-tat. It will be interesting in future empirical research to test whether these many examples of reciprocal cooperation may be based on learning combined with psychological preferences that value cooperation.

While our model shows that evolution can lead to intrinsically rewarding mutual cooperation, such utilities do not necessarily correspond to pure other-regarding preferences. For low benefit-to-cost ratios, competitive preferences that value other’s payoff negatively tend to evolve. In contrast, for sufficiently high benefit-to-cost ratio we see the evolution of conditional (multiplicative) other-regarding preferences, in agreement with previous results that found these preferences to be evolutionarily stable in continuous social dilemmas (Akçay et al. 2009). On the other hand, one could interpret our results for the freely evolving utilities as reflecting the evolution of the correct representation of real fitness effect of mutual cooperation, because mutual cooperation generates a positive effect on fitness. However, the “Realistic” utility function is not the only evolutionarily successful one in our model. For example, some evolutionarily successful preference functions value positively both mutual cooperation and mutual defection. These signs, together with a negative utility for the sucker’s outcome guarantee uninvadability by the Selfish preference function, because individuals with such preferences will learn to defect against Selfish. Moreover, on average the utility for the four different outcomes are ordered differently than the real material payoffs (for instance, mutual defection is the outcome with highest utility on average, while the real material payoff for this outcome is only the third material payoff). Therefore, our results do not show that natural selection leads to the correct representation of fitness effects in the brain in the context of learning. Another important distinction is that, even though the utility for the temptation to defect is the highest in the model with freely evolving utilities, this does not necessarily mean that there are no other-regarding preferences: in our simulations where the utility is a function of payoffs, other-regard (e.g., a positive *β*) can evolve even if the values of other evolutionary parameters make the temptation outcome being more rewarding than mutual cooperation.

Our finding that reward representations in the brain do not necessarily correspond to real fitness effects adds to a growing realization that natural selection can shape decision-making mechanisms to have specific biases in different ecological situations (e.g., McNamara et al. 2013). Generally, perceptual systems that represent the world accurately may not be evolutionarily stable, contrary to a naïve understanding of the workings of natural selection (Mark et al. 2010). In our scenario, the mismatch occurs because the game is repeated while an individual represents in his mind only the one-shot version of the game. It will be interesting in the future to examine whether our results obtained for the Prisoner’s dilemma extend to other repeated games.

### Diversity in preferences

Another main finding from our model is that a diversity of utility functions can be evolutionarily favored. This result is consistent with empirical findings that humans in behavioral experiments show behavioral diversity. In particular, strategies that value cooperation positively can produce similar behavior to that of reciprocating strategies (repeating the action of the partner in the previous round), and Selfish can produce the behavior of non-cooperators; these two behavioral types have recently been found to represent the action sequence of many human participants in laboratory experiments (Burton-Chellew et al. 2016; Fischbacher et al. 2001) and have been considered as plausible evolutionarily significant behavioral rules in theoretical models (Trivers 1971; Axelrod and Hamilton 1981; Lehmann and Keller 2006; Stewart and Plotkin 2013). Moreover, in addition to a diversity of preference types, our model also shows the potential for multiple behavioral outcomes in a population monomorphic for a given preference function. This is because of the fact that stochastic learning processes can converge to different equilibrium profiles, which provides another potential explanation for the behavioral variation observed in learning experiments (Chmura et al. 2012). That a single decision rule can produce behavioral polymorphism is a result that has been previously obtained in other models focusing on the evolution of cognitive mechanisms (Dridi and Lehmann 2015). This result illustrates that by modeling the evolution of the decision rules rather than the behavior themselves (McNamara and Houston 2009; Hammerstein and Stevens 2012; Fawcett et al. 2013; Dridi and Lehmann 2014), one can account for richer behavioral patterns and potentially provide insights into the psychological underpinnings of social behavior. Our model can indeed be viewed as capturing variations in the decision rules since by changing the utility function of an individual, the updating rule for action values also changes (see eq. 1), which subsequently produces different behavioral dynamics. These types of models require a detailed integration of two timescales (behavioral and evolutionary dynamics) and are consequently more difficult to analyze, but this difficulty cannot be avoided in trying to represent more realistically animal behavior.

In conclusion, our model articulates four levels of determinants of behavior: (1) the biological rewards at the core of brain functioning; (2) the psychological preferences that determine which states of the world are rewarding; (3) the social interactions that affect changes in the states of the world; (4) the biological process of natural selection determining which behavioral mechanisms prevail in an evolving population. We find that evolution of rewards for learning captures both the possibility of cooperation and a diversity of individual preferences that can be evolutionarily successful. These results show the promise of integrating learning based on evolving intrinsic rewards from social interactions as a proximate mechanism for understanding the nature of cooperation in humans and other animals.

## Appendix A Model details and analysis

### Behavioral analysis

In this section, we show how we analyze behavioral interactions by focusing on one particular example, Realistic vs. Other-regard (a Sagemath notebook that contains the analysis of all behavioral interactions is available on demand).

Treating Realistic as player *u* and Other-regard as player *υ*, one starts by verifying what behavioral equilibria of eq. 3 (Fig. B1) exist for this particular interaction (see Fig. B2 for the vector field of this interaction). Recall that the utilities of Realistic have the signs *u*_11_ > 0, *u*_12_ < 0, *u*_21_ > 0, *u*_22_ = 0. The utilities of Other-regard have the signs *υ*_11_ > 0, *υ*_12_ > 0, *υ*_21_ < 0, *υ*_22_ < 0. Consequently, the equilibria that do not exist are the two interior equilibria as well as 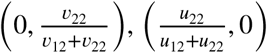, and 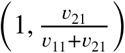. For example, the latter equilibrium does not exist because 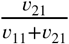 is either negative (when |*υ*_11_| > |*υ*_21_|) or greater than 1 (when |*υ*_11_| < |*υ*_21_|), which is impossible for a probability.

We then calculate the Jacobian matrix associated to eq. 3, evaluate it at each equilibrium, and calculate its eigenvalues. The pure equilibria (0, 0), (1, 0), (0, 1), and (1, 1) are straightforward to analyze because the sign of the eigenvalues are opposite to the sign of the utilities of the players. For example, the equilibrium (1, 1) has the associated eigenvalues (−*ξu*_11_, −*ξυ*_11_), which makes it locally stable. The equilibrium (0,1) is also locally stable because it has the associated eigenvalues (−*ξu*_21_, −*ξυ*_12_), which are both negative. The equilibria where one player is mixing require a little more work. For example, the equilibrium 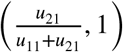 has eigenvalues

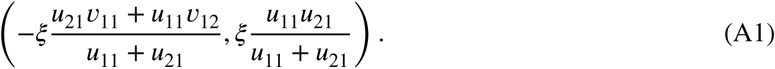

Solving the inequality

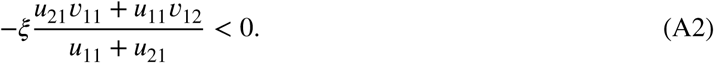
shows that this is always true given the signs of **u** and **v**. The second eigenvalue

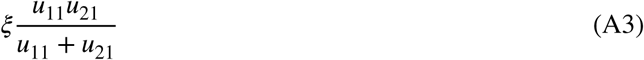
is always positive, making the equilibrium 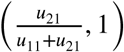 a saddle.

### Fitness computation

In this Appendix, we delineate the logic behind the computation of the fitnesses in Table 1 for the interaction between Realistic and Other-regard. The other fitnesses are computed similarly.

As we have proven in the section “Behavioral analysis”, the interaction between Realistic and Other-regard can lead to two possible behavioral equilibria, (1, 1) or (0, 1). Because the original learning dynamics is stochastic, it may reach either equilibrium, but we cannot determine which one analytically. Indeed, the theory of stochastic approximations does not provide precise predictions about the lock-in probabilities in local attractors. Hence, in the analysis of the model we treat the probability *q*_RO_ to get attracted in the mutual cooperation equilibrium, (1, 1), as a free parameter. Hence the probability to reach the equilibrium (0, 1) is 1 − *q*_RO_. The payoff to the players at equilibrium (1, 1) is *b* − *c* for both players. At equilibrium (0, 1), Realistic obtains *b* and Other-regard −*c*. Weighting these payoffs by the lock-in probabilities gives

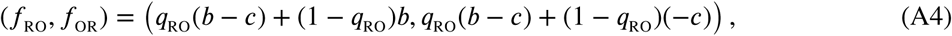
which are the entries of the corresponding fitnesses in Table 1 of the main text.

### Replicator dynamics and evolutionary equilibria

We evaluate the evolutionary competition between Realistic, Other-regard, Selfish, and Manipulator using the 4-dimensional replicator dynamics. In order to do so, we need to construct the evolutionary fitness matrix, i.e.,computing the fitness, *f*_*uυ*_, of a strategy *u* when matched with another strategy *υ* in the repeated game. We need to do so for all strategies *u, υ* ∈ {R, O, M, S}, where we denote a strategy by its initial (i.e., R denotes Realistic, etc.). Having obtained such fitnesses, one can then call **x** = (*x*_R_, *x*_O_, *x*_M_, *x*_S_) the vector of frequencies in the population (*x*_R_ + *x*_O_ + *x*_M_ + *x*_S_ = 1) and define the average fitness of type *u* when the population is in state **x** as

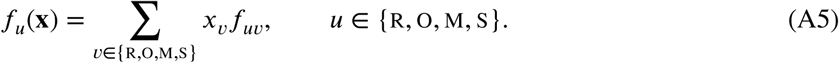

We then use the continuous-time replicator dynamics to assess the long-term frequencies of strategies in the population, i.e.,

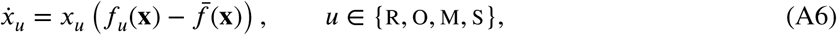
where

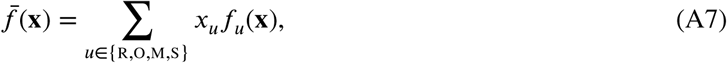
is the average fitness in the population at state **x**.

Solving for the equilibria of eq. A6, we find that there is an equilibrium with Realistic, Other-regard, and Selfish, given by

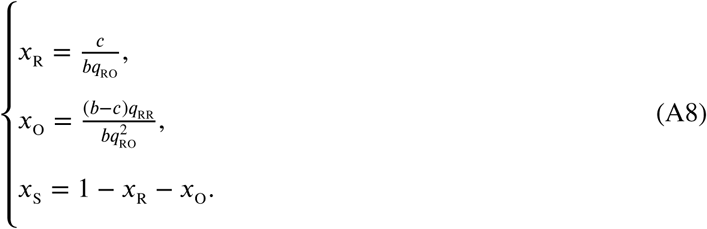

We find another 3-strategy equilibrium with Realistic, Manipulator, and Selfish, with the frequencies

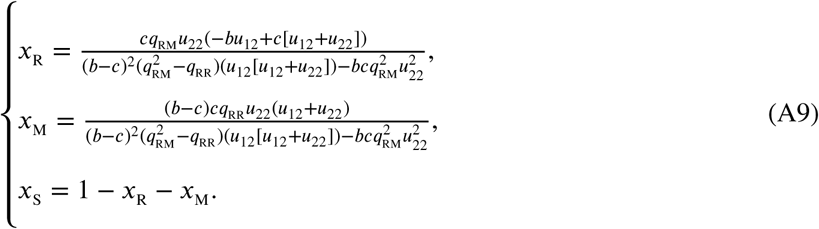

There exists a 2-strategy equilibrium with Realistic and Other-regard who are in frequencies

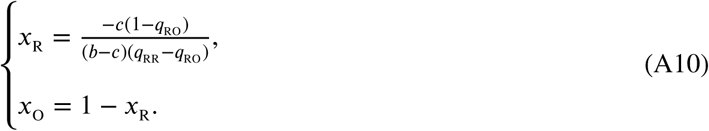

Selfish invades the latter mix of Realistic and Other-regard, and makes part of the equilibrium when

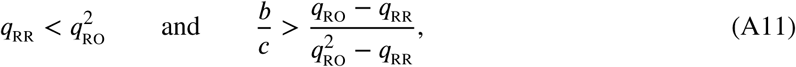
which is obtained by finding the conditions under which the fitness of Selfish at the equilibrium of eq. A10, 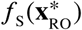, is higher than the average fitness in the population, 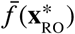. When this occurs the equilibrium consisting of Realistic, Other-regard, and Selfish (eq. A8) becomes stable.

## Appendix B Supplementary Tables and Figures

**Table B1:** Classification of behavioral outcomes amongst the 4 strategies considered.

| Interaction | Stable equilibria |
| --- | --- |
| R vs. R | (1, 1) and (0, 0) |
| R vs. O | (0, 1) and (1, 1) |
| R vs. M | (1, 1) and $(0, \frac{v_{22}}{v_{12}+v_{22}})$ |
| R vs. S | (0, 0) |
| O vs. O | (1, 1) |
| O vs. M | (1, 0) and (1, 1) |
| O vs. S | (1, 0) |
| M vs. M | (1, 1)* |
| M vs. S | $(\frac{u_{22}}{u_{12}+u_{22}}, 0)$ |
| S vs. S | (0, 0) |

\*for $|u_{12}| < |u_{21}|$ , another equilibrium, $(\frac{u_{22}}{u_{12}+u_{22}}, 0)$ , becomes stable but we place ourselves in the conditions where this condition does not hold.

**Figure B1:**
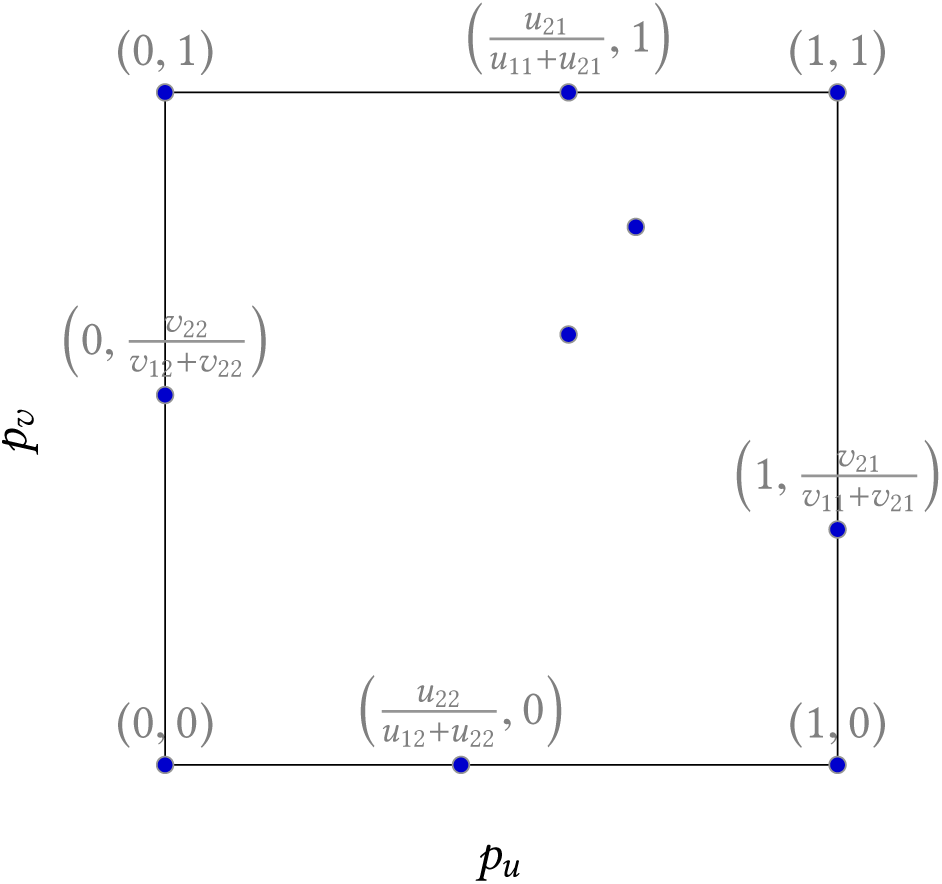
The ten generic behavioral equilibria in a 2 × 2 game between a player with utility function *u*, and his opponent with utility function *υ*. The two interior equilibria have long expressions that are not shown here.

**Figure B2:**
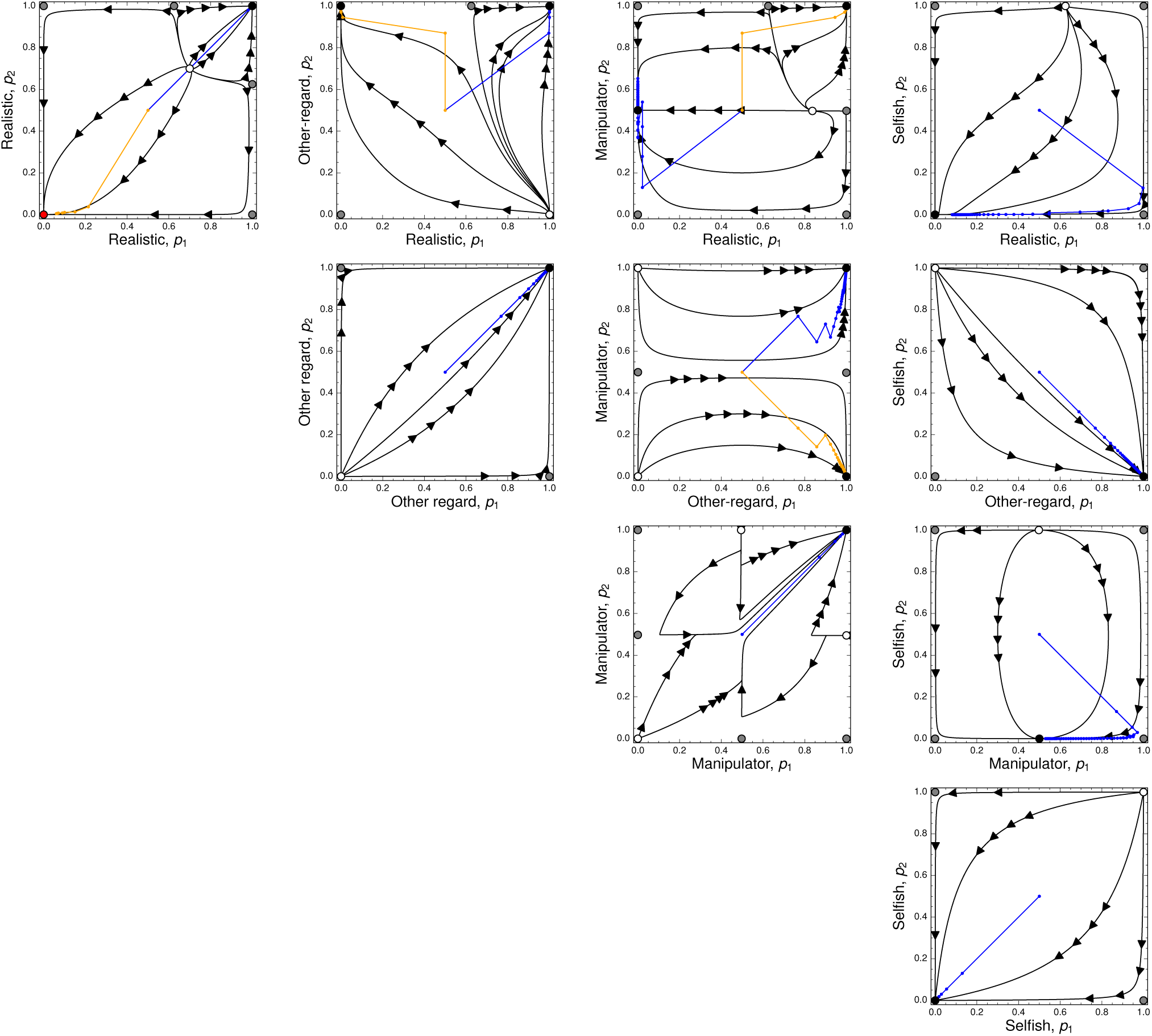
Solution trajectories (black) and stochastic trajectories (colored lines) for the 10 possible behavioral interactions between the 4 strategies Realistic, Other-regard, Manipulator, and Selfish. In each panel, on the *x*-axis is represented the probability that the row player cooperates (*p*_1_), while on the *y*-axis, this is the probability that the column player cooperates (*p*_2_). The stochastic trajectories are started from the center of the state space 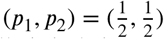 and dots on it represent interaction rounds between the players. Circles represent equilibria: a white-filled circle is a source (both associated eigenvalues are positive); a gray-filled circle is a saddle (one positive and one negative associated eigenvalue); a black circle is a sink (both associated eigenvalues are negative). The red circle in the Realistic-Realistic interaction is a degenerate equilibrium with both zero eigenvalues, but it turns out to be locally stable. These plots were generated by generally setting positive utilities close to 1 and negative utilities close to −1 (which is why mixed equilibria always appear close to 0.5) except for the Realistic strategy for which we set the utility matrix to equate the payoff matrix of the Prisoner’s dilemma with *b* = 5 and *c* = 3.

**Figure B3:**
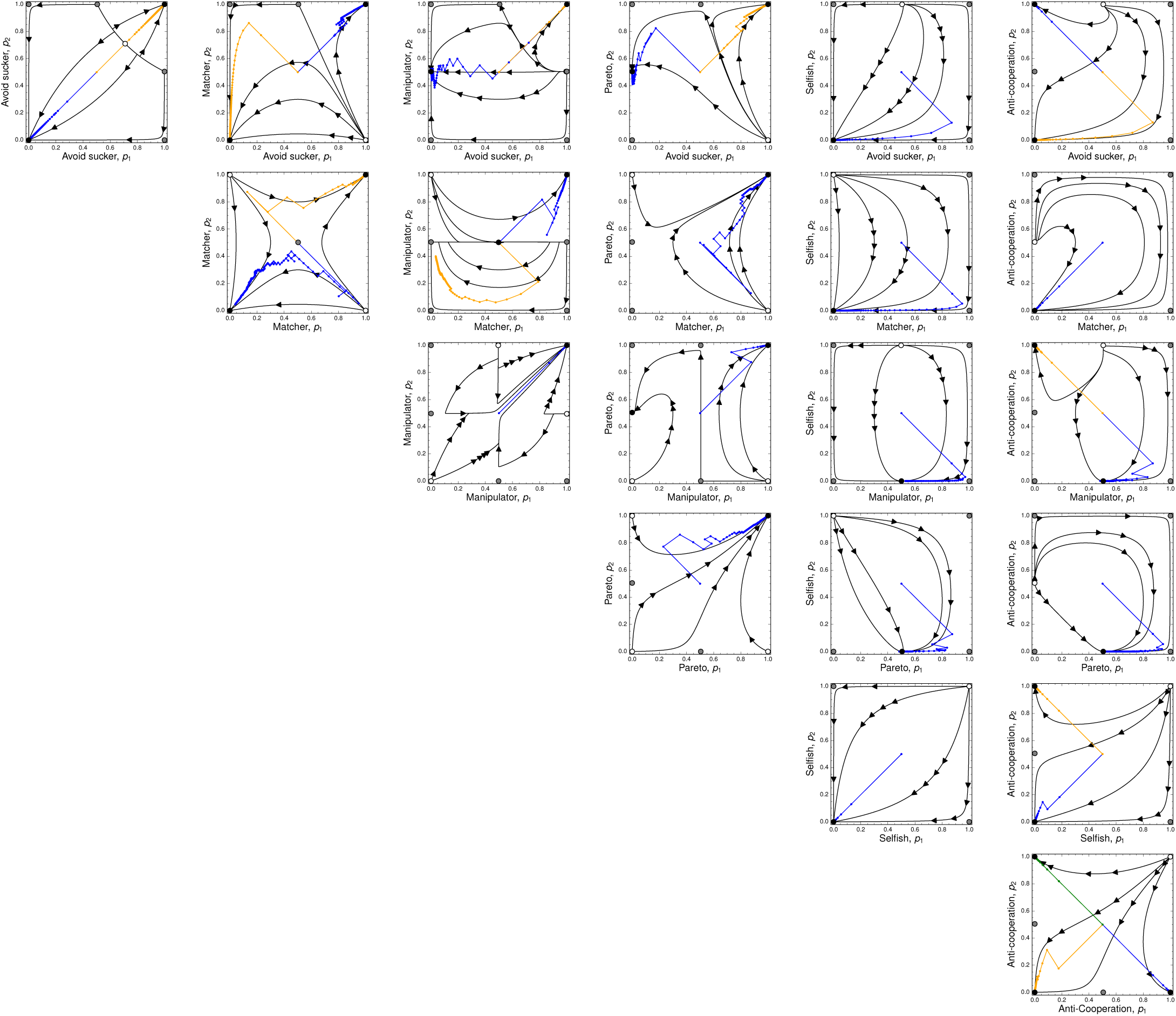
Solution trajectories (black) and stochastic trajectories (colored lines) for the 21 possible behavioral interactions between the 6 dominant strategies in the simulations: Avoid Sucker’s Payoff, Matcher, Manipulator, Pareto, Selfish, and Anti-Cooperation. Otherwise similar to Fig. B2.

**Figure B4:**
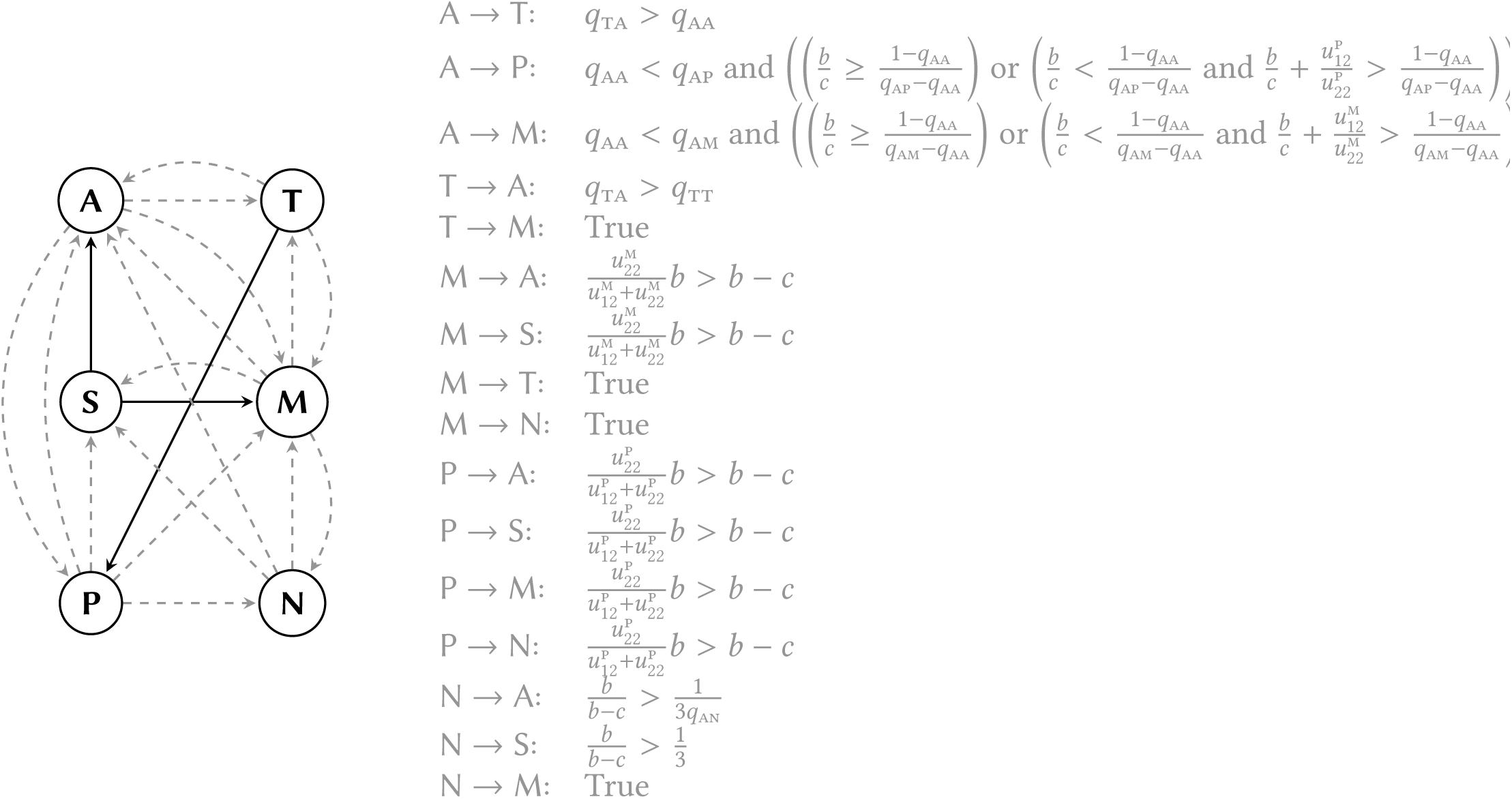
Invasion diagram amongst the successful strategy classes in the evolutionary simulations with associated invasion conditions (similar to Fig. 2B). When we write “True”, the invasion condition exists but is too long to be displayed here (a Mathematica notebook containing these conditions is available on demand). Name legend: A=“Avoid Sucker’s Payoff”; T=“Matcher”; P=“Pareto”; N=“Anti-cooperation”; S=“Selfish”; M=“Manipulator”.

**Figure B5:**
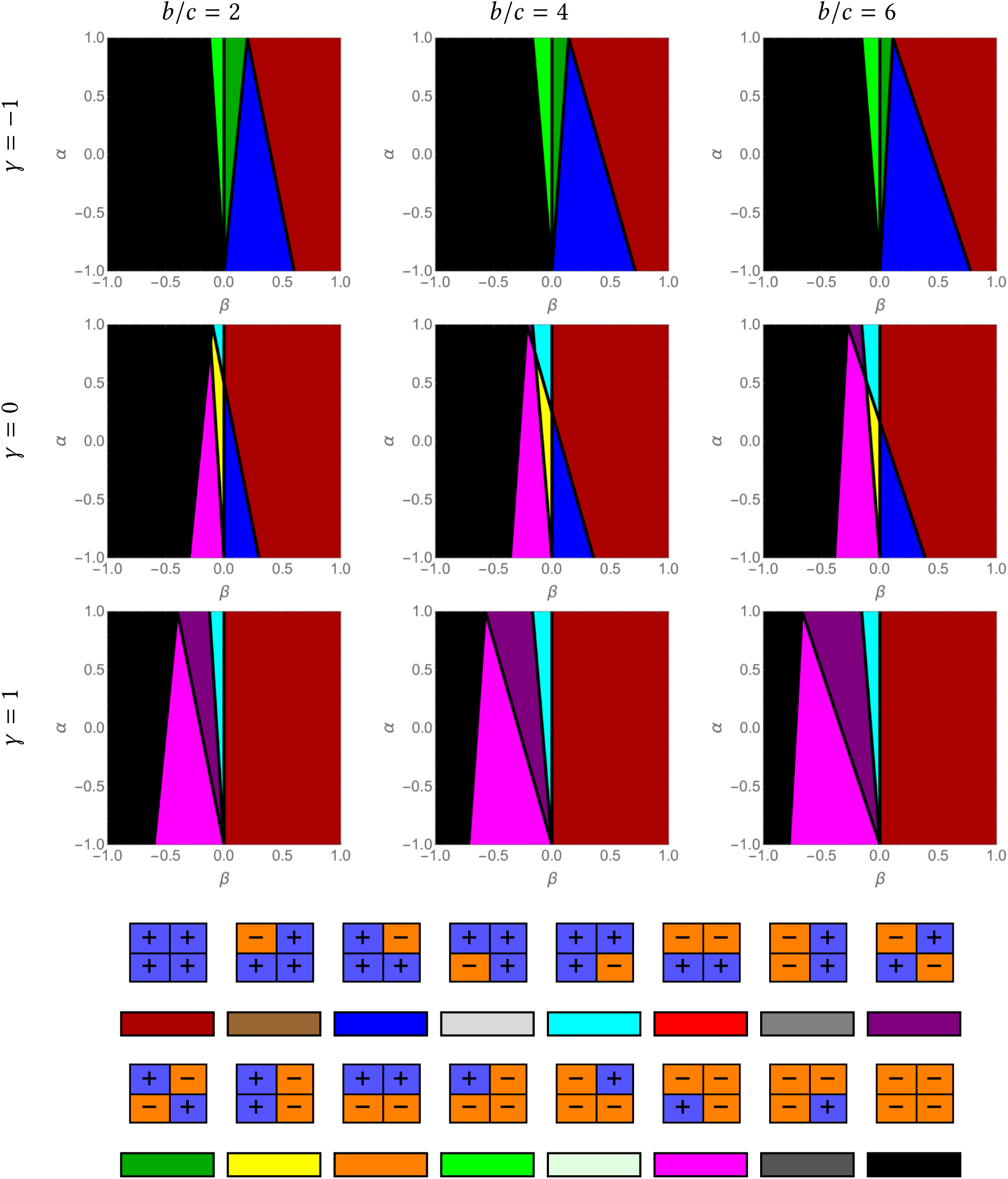
Phenotype space for the model where the utility depends explictly on material payoffs (eq. 6 with *k* = 2) as a function of *β* (*x*-axis), *α* (*y*-axis), *γ* (rows), and *b*/*c* (columns). Under the panels depicting the phenotype space, we show the color code of strategies (colors are under the corresponding utility matrix).

**Figure B6:**
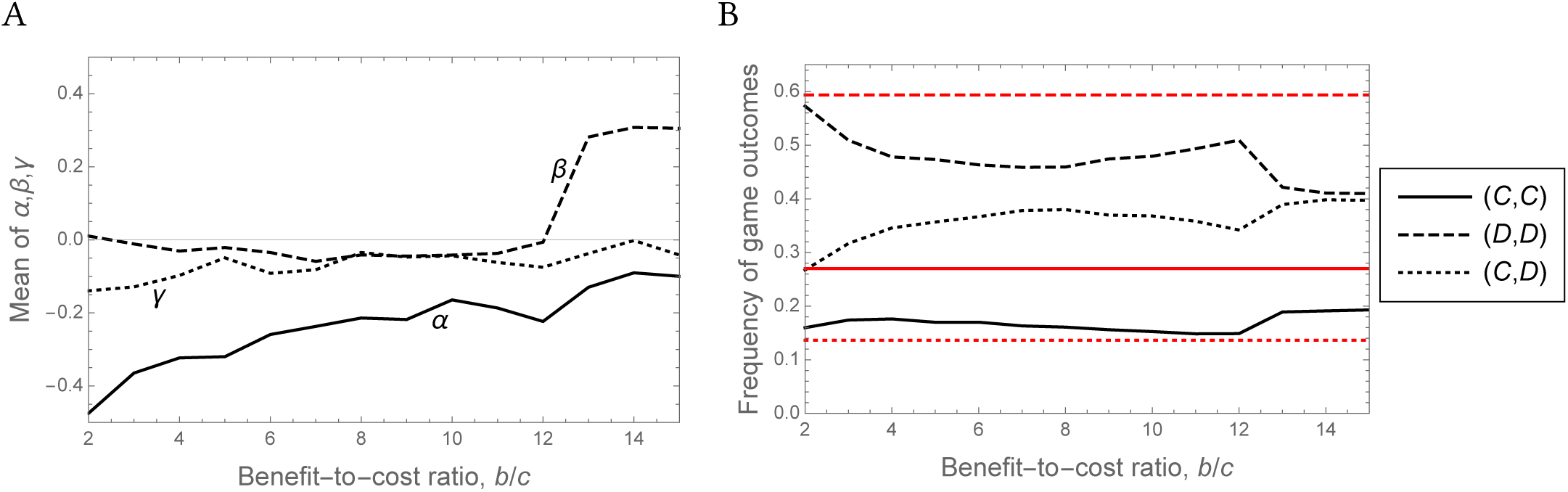
Results for the model where the utility depends explictly on material payoffs (eq. 6) as a function of the benefit-to-cost ratio, *b*/*c* (identical to Fig. 5 except that *k* = 5). (A) Time average of *α, β, γ* in a simulation run. (B) Time average of the frequency of game outcomes in a simulation run (similar to Fig. 4B and 5B)

**Figure B7:**
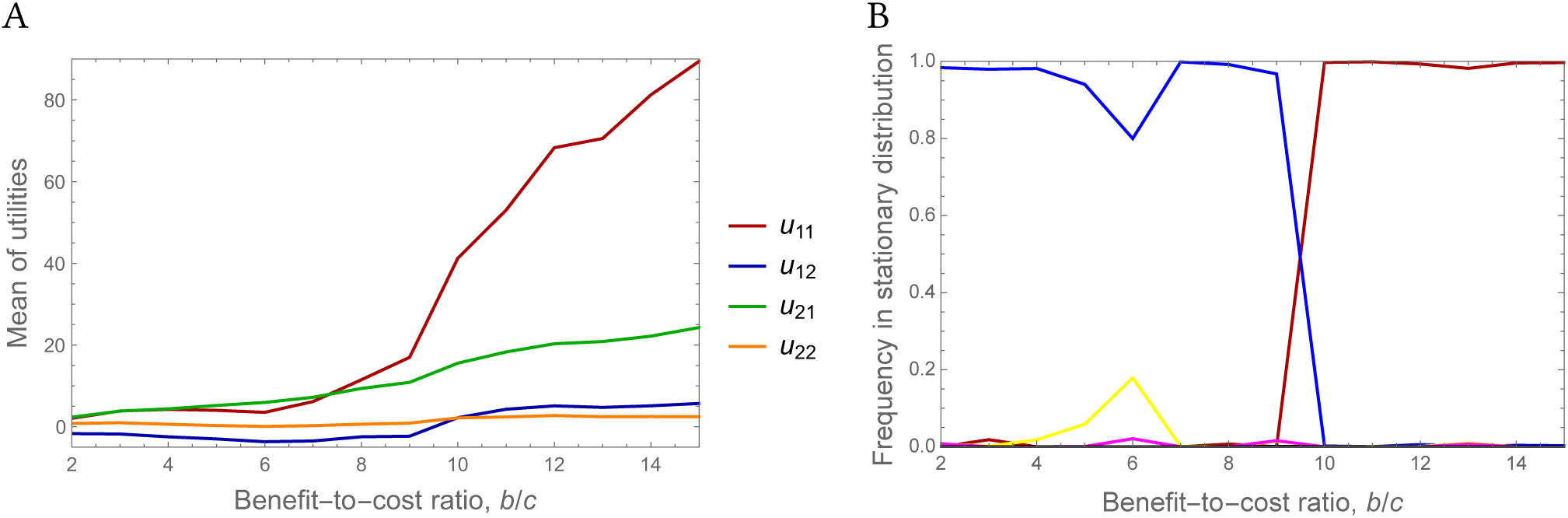
Additional results for the model where the utility depends explictly on material payoffs (eq. 6 with *k* = 2) as a function of the benefit-to-cost ratio, *b*/*c*. (A) Time average in a simulation run of the four utilities associated to each game outcome. (B) Proportion of time a simulation run spends in each strategy class (similar to Fig. 4A; see Fig. B5 for color code of strategies).

**Figure B8:**
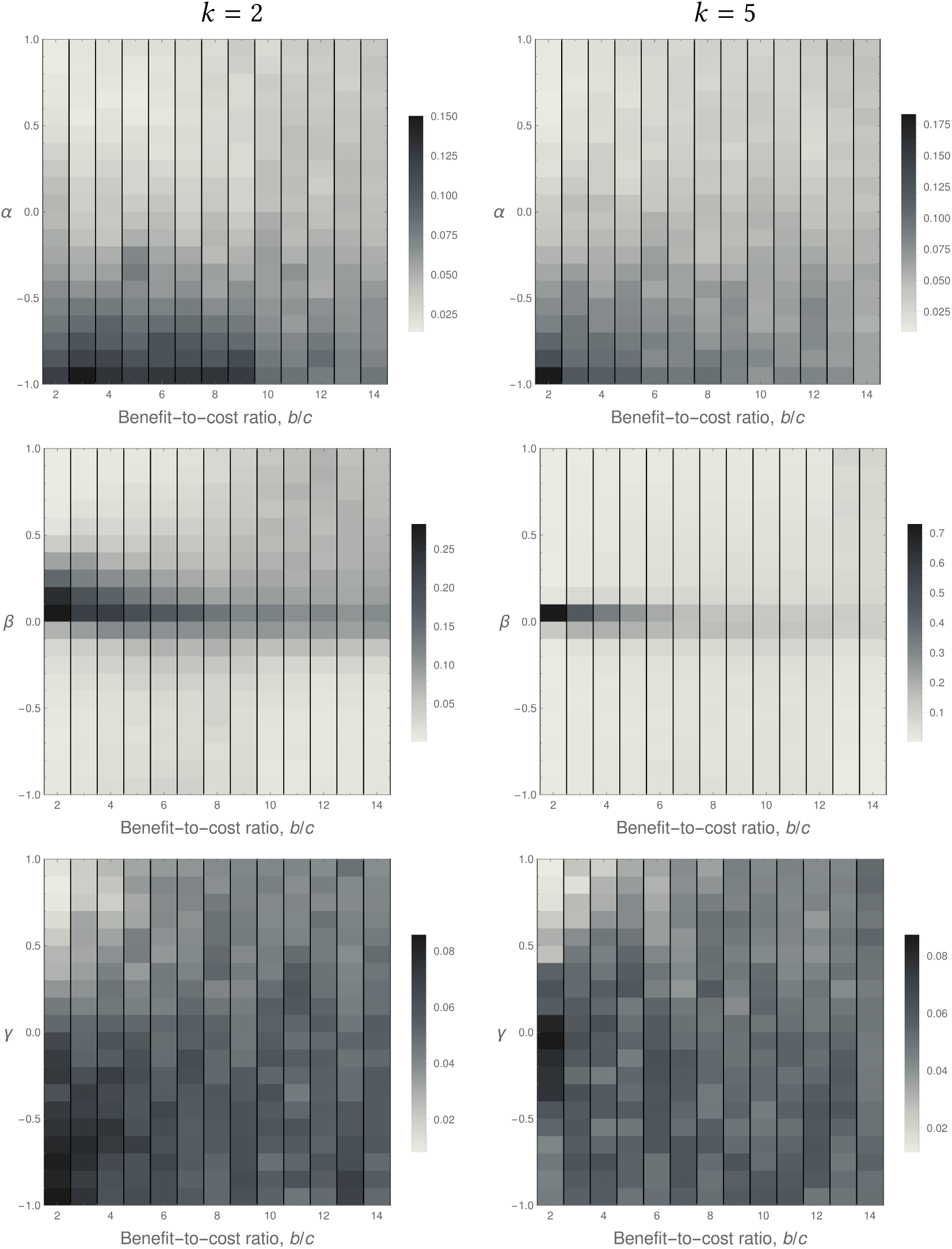
Stationary distributions of the traits measuring additive other-regard (*α*), multiplicative other-regard (*β*), and inequity aversion (*γ*) for various benefit-to-cost ratios (*b*/*c*) and values of *k* (see main text for definition, eq. 6) when constraining the utility function to explicitly depend on the material payoffs. In each subfigure, a column of values for a given benefit-to-cost ratio represents the stationary distribution of the trait, where darker shading corresponds to a higher frequency.

**Figure B9:**
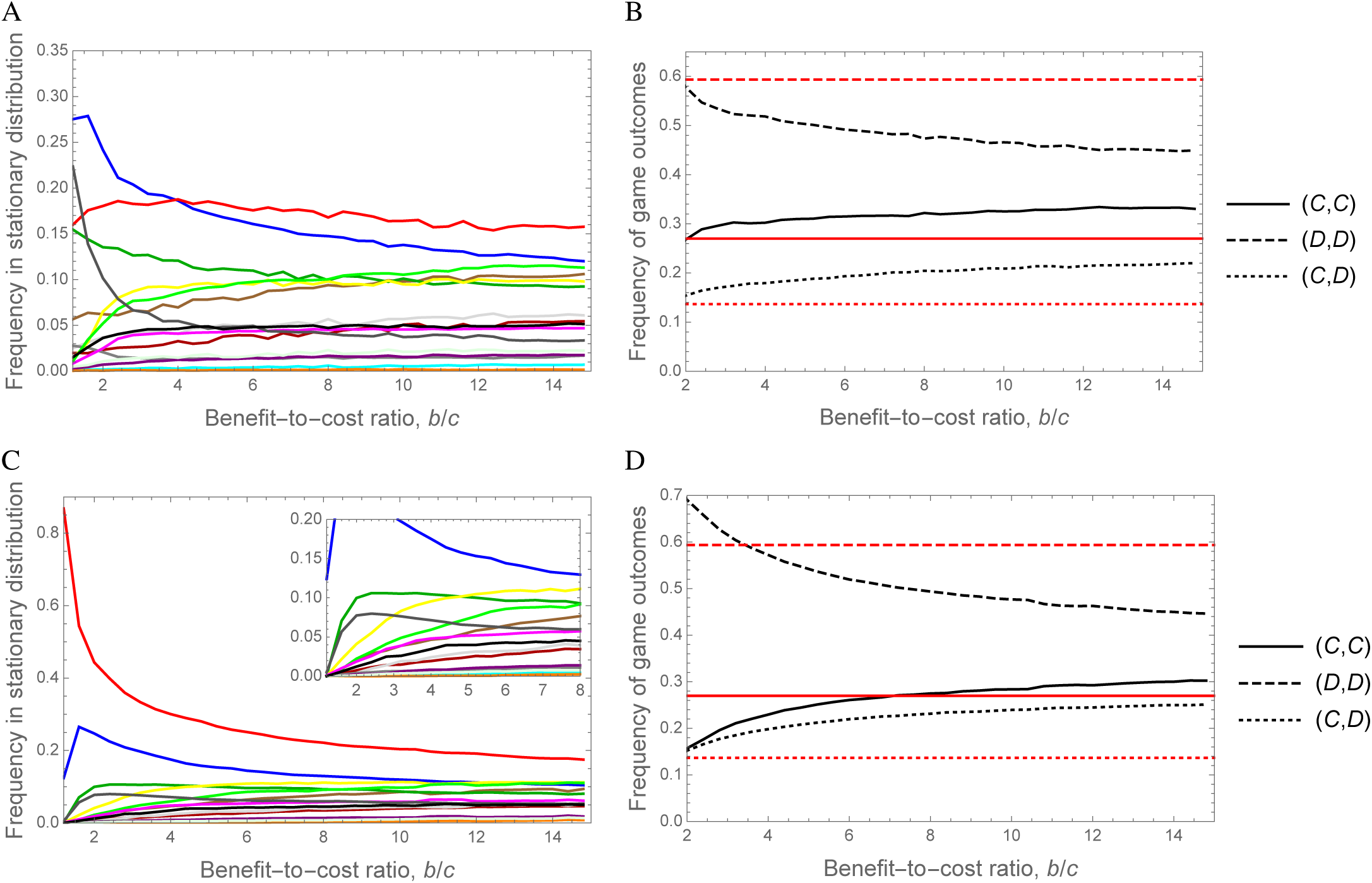
Reproduction of Fig. 4 for lower values of the game duration, *T*. A and B: *T* = 50; C and D: *T* = 10. A and C: Proportion of time the simulation spends in each of the 16 strategy classes as a function of *b*/*c*. Colors of strategies are as in Fig. 3 (see also Fig. B5). B and D: Time average of the frequency of game outcomes in a simulation run as a function of *b*/*c* (black lines). The red lines show the expected frequency of the corresponding game outcomes in the population if we draw strategies randomly from a uniform distribution. Other parameter values than *T* are as in Fig. 4

**Figure B10:**
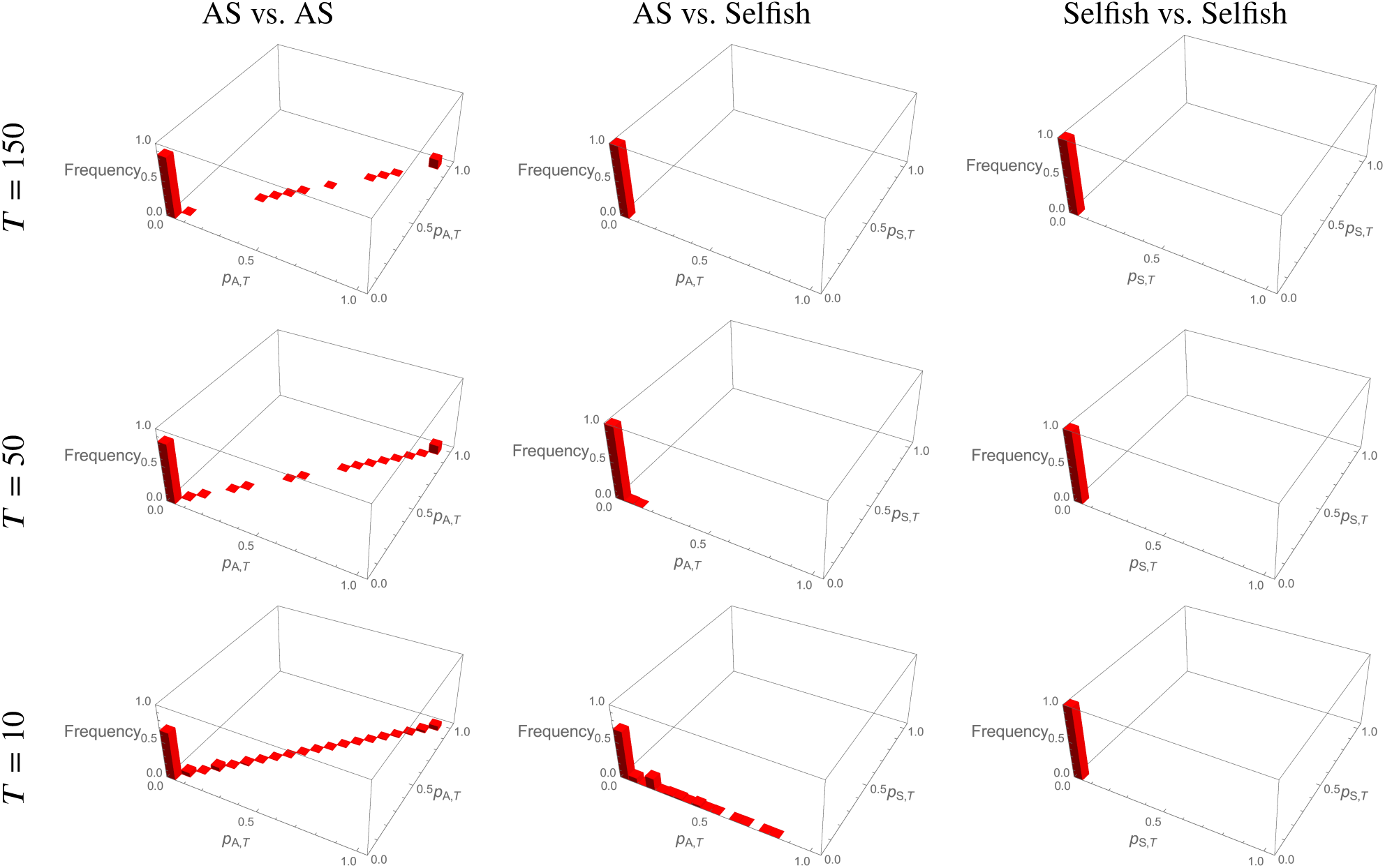
Distribution of behavior at the end of the game, time *T*, for matches involving the “Avoid Sucker’s payoff” strategy (AS) and Selfish. For each possible type of pairing and duration of the game *T*, we simulate learning between *N* = 3000 players (i.e., 1500 pairs), and we show here the frequency of pairs displaying a particular combination of probabilities to cooperate, *p*_*i,T*_. Other parameter values are as in Fig. 4.

